# Genome-wide differentiation and SNP-based identification of northeastern Atlantic *Sebastes* species

**DOI:** 10.1101/2025.04.11.648347

**Authors:** Jansson Eeva, Besnier Francois, Christiansen Henrik, Saha Atal, Tranang Caroline Aas, Bruvold Ingrid Marie, Mateos Rivera Alejandro, Johansen Torild

## Abstract

Sustainable fisheries require reliable species identification and understanding of the underlying genetic hierarchies within the targeted species. Three *Sebastes* species are commonly found in the northeastern Atlantic: *Sebastes mentella* (beaked redfish), *Sebastes norvegicus* (golden redfish), and *Sebastes viviparus* (Norway redfish). These species are morphologically similar and have largely overlapping distribution ranges. Besides, three cryptic species for *S. norvegicus* and three depth-defined ecotypes for *S. mentella* have been suggested. Genetic knowledge and methods are needed to identify and monitor these species and to look at their geographic distribution.

Here, a total of 99 specimens of *S. mentella, S. viviparus* as well as *S. norvegicus* A and B were sequenced in pools and aligned against a reference genome from a sister species, *S. fasciatus* (Acadian redfish). The measured divergence between all pairs, including the cryptic species pair *S. norvegicus* A and B, was high (mean *F*_ST_ = 0.33-0.61) and encompassing throughout genomes. Several shared megabase-scale regions of elevated divergence were observed, likely representing regions of reduced recombination. Moreover, 2914 fish collected across the northeastern Atlantic were analysed with a discriminatory SNP panel of high resolution (mean *F*_ST_= 0.60-0.91), revealed few possible hybrids, and supported further sub-structuring within *mentella* and *S. norvegicus* B. The latter was split into two groups, one of which was the previously recognized ‘giant’ morph and confirmed here for the first time in Norway. *Sebastes mentella* had two to three groups, likely representing previously identified depth-related ecotypes.

Our study shows high genetic distinctiveness between acknowledged northeast-Atlantic *Sebastes* species, and between the cryptic species *S. norvegicus* A and B. Previously identified, fast-growing ‘giant’ seems to be closely related to *S. norvegicus* B. Only few SNP markers are necessary for accurate species determination, facilitating further studies and widely applicable monitoring.

## 1. Background

Marine resources are used and depleted at an accelerating pace [1]. Changing climate and multiple other, coincident anthropogenic stressors threaten many marine species. Concurrently, marine biodiversity is underprotected [2] and understudied [3]. Specifically, intraspecific, genetic diversity is often neglected in management and conservation actions [4, 5]. Species that resemble each other, or are morphologically indistinguishable (i.e. cryptic), pose a special conservation challenge [6]. For commercial species in particular, it is crucial to accurately understand and identify species boundaries, co-occurrence, hybridization, and simultaneous exploitation of closely related and/or cryptic species. Genetic methods offer means to not only better understand the marine biodiversity, but also to efficiently protect and sustainably exploit cryptic marine species [7, 8].

Redfish belong to the genus *Sebastes*, commonly also called rockfishes, that consists of over 110 described species with remarkable morphological and ecological diversity [9, 10]. These species are bound to temperate-cool, upwelling driven systems, and the majority of them live in the Pacific. Four species, *Sebastes mentella* (beaked redfish), *S. norvegicus* (golden redfish; previously misattributed to *S. marinus*), *S. viviparus* (Norway redfish), and *S. fasciatus* (Acadian redfish) inhabit North Atlantic waters [9]. The distribution of *S. fasciatus* is concentrated on the western Atlantic side [11], and *S. viviparus* is common on the Northeast Atlantic[12], whereas *S. mentella* and *S. norvegicus* are transatlantic [11, 13, 14]. These species are long-lived, grow slowly and mature late [15, 16] – all characteristics that make them vulnerable for overexploitation. Additionally, redfish produce relatively seldom strong year classes and at irregular intervals [16]. Atlantic redfish species have somewhat differing depth preferences but co-occur in many areas [11], and hybrids have been reported [17, 18]. Together with modelled secondary contacts [11], this suggest that barriers between the species are not absolute.

North Atlantic *Sebastes* species diverged relatively recently, ∼3 MYA [9], and exhibit subtle morphological differences so that their distinction using traditional methods can be tricky [19]. All of these species are commercially exploited, albeit *S. viviparus* at low levels due to its smaller size [20]. Mixed fisheries and possible (unintentional) depletion of species, specific ecotypes and/or genetically different populations is of high management concern [16, 21]. Genetic approaches can provide important management information [22] including knowledge of underlying population structure, hybridization, and adaptive variability [23]. North Atlantic *Sebastes* species have been studied genetically since early 1990s, starting with few allozymes (e.g.[12, 19]) to recent investigations, where thousands of markers [11, 14, 24] or even entire genomes are screened. In general, more markers and better genomic coverage add resolution, and enable studies of e.g. adaptive divergence [23]. The acknowledged four North Atlantic *Sebastes* species are genetically distinct, but additionally, *S. mentella* has been shown to consist of at least three genetically diverged depth-related ecotypes (shallow pelagic, deep pelagic, and demersal slope; [11, 14, 18, 25]), *S. norvegicus* likely includes three cryptic species (named as A, B, and giant; [13, 26] see also [27]), and that *S. fasciatus* has at least five highly diverged populations [11]. Now, with the first chromosome level reference genome for a North Atlantic *Sebastes* species, *S. fasciatus*, available (https://www.ncbi.nlm.nih.gov/datasets/genome/GCA_043250625.1/), differences among (and within) species can be anchored into a genomic landscape [28], enabling many new research possibilities.

Our aim in this study was two-fold: I) to describe the amount and general pattern of genome-wide divergence between *Sebastes mentella*, *S. norvegicus* A and B, and *S. viviparus*, and II) to test the performance of the recently published [24] SNP panel for *Sebastes* species identification across the northeastern Atlantic (from the Barents Sea and Norwegian coast in the East to west Greenland coast in the West). Here, we specifically aim to document redfish species composition in different areas, to identify possible hybrids, and to test whether the SNP panel can discriminate the substructures mentioned above, i.e. cryptic species and ecotypes.

## 2. Material and methods

This study consists of two intervened parts. First, fish of each target species – *Sebastes mentella*, *S. norvegicus* A, *S. norvegicus* B, and *S. viviparus* – were collected and sequenced in pools [29], hereafter referred to as “pool-seq” to characterize the general genomic divergence patterns between the species. This dataset was also used to develop species-specific SNP panels (as explained in detail in Johansen *et al*. 2025). In the second part of this study, the performance of these SNP panels was scrutinized on a large collection of *Sebastes* specimens (2914 individuals) throughout the northeastern Atlantic distribution area.

### 2.1 Sampling

For pool-seq, a total of 99 fish were used (STable 1). Fish were collected between 2001 – 2016 in Norway, Greenland, and the Faroe Islands. From 16 to 44 fish per species were used per pool.

For the SNP panels, samples were collected between 2016 – 2023 from *Sebastes* occurring in the Northeast Atlantic, i.e. around Iceland and East from Greenland (Figure 1), and with special focus on Norway’s exclusive economic zone. Norwegian coastal and shelf areas were collected repeatedly as part of annual research cruises (STable 2), whereas the remainder were from surveys, commercial fisheries, or unique sampling occasions like the four specimens caught in Skagerrak by sport fishermen. Sampling was opportunistic, and sampling coverage therefore highly uneven. For most specimens, assumed species was recorded upon catch by trained experts or experienced fishermen together with geographic coordinates. Most samples were analysed for morphological characteristics (see Bruvold *et al.* (2025)). To guarantee consistency, we included known reference samples of *S. norvegicus ‘*giants’ and *S. norvegicus* A, as presented by Saha *et al.* (2017).

**Figure 1.**
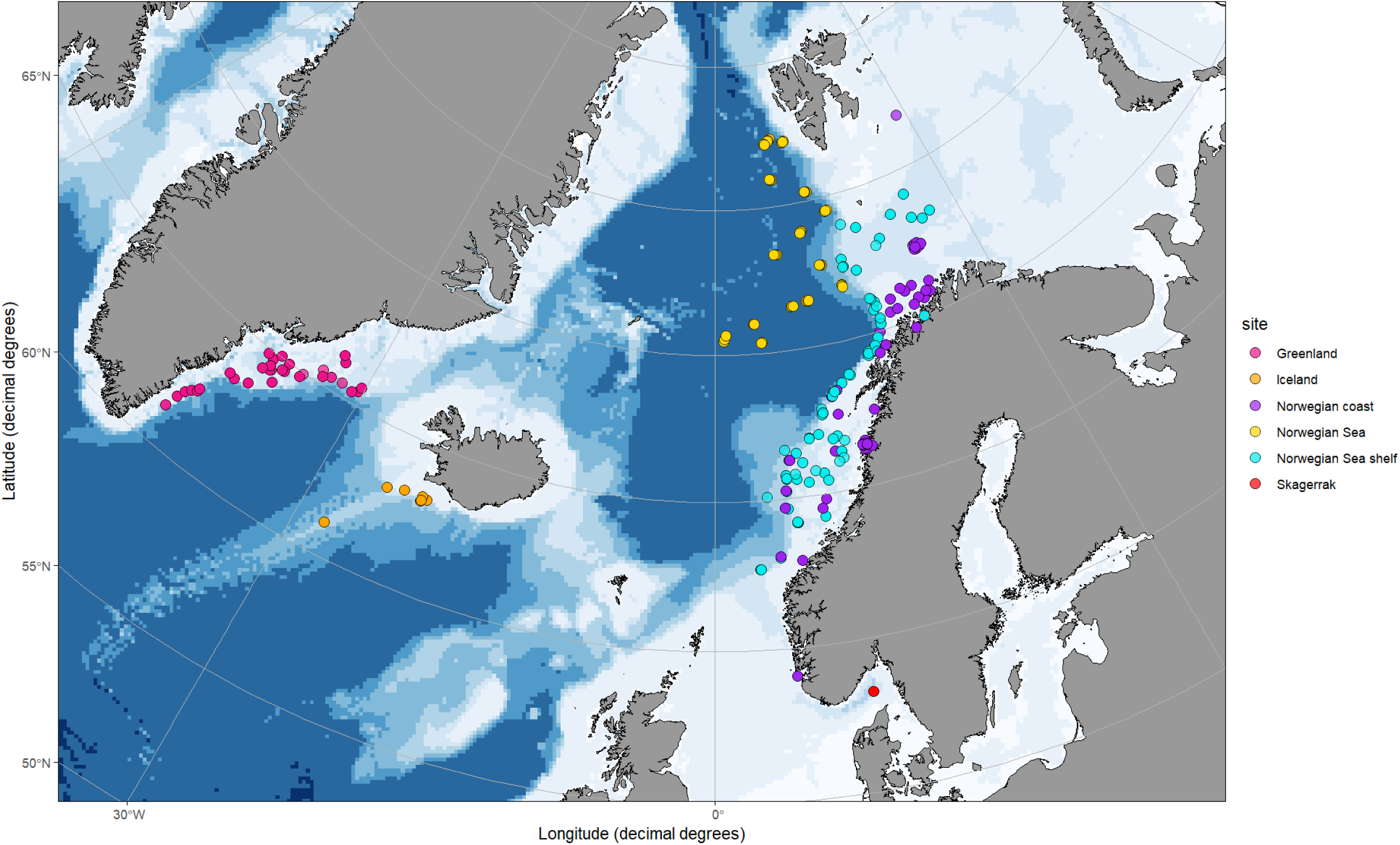
Map of sampling regions covered in this study. Given points indicate individual specimens. Please note that these groupings were not used in any analysis but set for illustrative purposes only. Details of the collected samples are given in STable 2.

### 2.2 DNA isolation

Fin clips of each fish were stored in absolute ethanol upon catch and used as a starting material. For the SNP panels, DNA was isolated using the E-Z 96 Tissue omega DNA Kit (Omega Bio-Teck, Inc.) following the manufacturer’s protocol. DNA from samples for pool-seq were isolated with Qiagen blood and tissue kit following the manufacturer’s protocol.

### 2.3 Pool-seq, quality control and SNP calling

DNA quality was checked visually on agarose gel. DNA with high molecular weight from all specimens was quantified using fluorometry (Qubit; Thermo Scientific) and pooled at equal concentrations, ∼20ng/µl, of genomic DNA for sequencing. The Oslo University Hospital in Oslo, Norway, conducted the construction of paired-end libraries followed by sequencing using read length of 150 bp on Illumina HiSeq 2500 machine.

Filtering for adaptor and overrepresented sequences was performed with *fastp* (https://doi.org/10.1093/bioinformatics/bty560). Recently published reference genome of closely related *S. fasciatus* (GCA_043250625.1; https://www.ncbi.nlm.nih.gov/datasets/genome/GCA_043250625.1/) was used as a reference against which pool-seq data was aligned with *bwa-mem2* [7], followed by SNP calling with *samtools* v.1.8 [8]. The R package vcfR [30] was used to visualise the coverage and mapping quality across all SNPs. SNPs were then filtered using *bcftool* together with custom R script to match the following criterion: minimum coverage = 50, maximum coverage = 250, minimum phred scaled quality (QUAL) = 200.

### 2.4 SNP panel development and selection

SNP panel development is described in detail in Johansen *et al.* (2025). In brief, sequencing data was aligned against a scaffold-level reference genome of *S. norvegicus* (ASM90030265v1; https://www.ncbi.nlm.nih.gov/datasets/genome/GCA_900302655.1) and screened for the most differentiated SNPs based on measured *F*_ST_ between each species pair. Primer design, amplification and genotype calling were conducted using the Agena MassARRAY iPLEX Platform as described by Gabriel *et al*. [31]. The final SNP selection consisted of 64 SNP markers passing all the quality filters explained above. Markers were arranged into three assay panels with ∼25 SNPs in each. Samples were processed on 384-plates including negative controls. Two panels were designed to separate different redfish species (ID panel); first, to distinguish *S. mentella, S. viviparus* and *S. norvegicus*, and second, to separate the two known cryptic *S. norvegicus* species, A and B [13]. The third panel was added to include population structure screening within *S. mentella* [18]. The development of these markers is described in Saha *et al*. [14].

### 2.5 Genotyping and quality control

All collected 2914 fish were genotyped for the two ID panels, and 1990 of them for the additional third panel. All data processing and analyses were done with *R* v. 4.1.2 [32] and its associated packages unless otherwise stated. Data manipulation and cleaning was done using *dplyr* v. 1.1.4 [33] and *janitor* v. 2.2.0 [34] packages, and visualized with *ggplot2* v. 3.5.0 [35]. The two datasets were quality checked separately. We included only loci that produced non-ambiguous clustering patterns (i.e. three clear, separable genotypes) on the used genotyping platform. Subsequently, R package *dartR* v. 2.9.7 [36, 37] was used to remove loci with more than 20% missing data as well as individuals with more than 30% missing data. Duplicated genotypes were removed.

### 2.6 Analysis of pool-seq data

All analyses of the pool-seq data were based on a set of SNPs that satisfied the quality criterion of minimum coverage =50, maximum coverage =250, minimum phred scaled quality (QUAL) =200. Using *Poolfstat* R package v. 3.0.0 [38], F-statistics were calculated between pairwise samples for the entire genome and in consecutive sliding windows of 100 SNPs along the 24 chromosomes present in the reference genome. Similarly, Principal Component Analysis (PCA) were obtained using the *randomallele.pca()* function in *poolfstat* package.

### 2.7 Genetic analysis of the SNP panel data

For most samples, assumed species had been recorded upon catch based on fish appearance or during a more detailed morphological assessment. This knowledge was utilized in the initial grouping phase explained below. After fish were grouped according to their likely species, genetic characteristics were investigated species-wise as well as within each species.

#### 2.7.1 Grouping of samples into assumed species based on principal components

First, we employed an assumption-free clustering approach, principal component analysis (PCA) using R package *ade4* v. 1.7-22 [39, 40]. Individual coordinates for all fish were retrieved from the produced PCA plot with *factoextra* v.1.0.7 [41], and observed clusters assigned to probable species using prior knowledge of species ID based on morphological/field identification. The same software was used to assess individual locus contributions, i.e. their resolution on the first two axes. To check for consistency, grouping was done also with *dapc* approach implemented in R package *adegenet* v.2.1.10 [42, 43]. Here, we first defined the most probable number of genetic groups (*K*) with the function *optim.a.score()* after which this number was used to assign individuals into their groups with *find.clusters.genind()* function. Concurrently, the division of species within each sampling location was recorded.

#### 2.7.2 Genetic clustering and estimation of admixture proportions

Next, we used STRUCTURE v.2.3.4 [44, 45], a model-based Bayesian clustering method that uses a predefined number of *K* clusters to assess the posterior probabilities of each individual’s genotype to originate from each cluster. We performed runs using the default admixture model with correlated allele frequencies, 20 000 *MCMC* (Markov Chain Monte Carlo) repeats as initial burn-in, followed by 50 000 repetitions in the actual run. We tested *K* values from 1 to 10, with three iterations for each value. To determine most likely clustering, we visually inspected obtained barplots together with *StructureSelector* software [46] that summarizes run results as the optimal *Ln Pr*(X|K) ad hoc summary statistic *ΔK* [47], which identifies the highest level of population hierarchy. In addition, the software produces statistics explained in Puechmaille [48] that adjust for uneven sampling between the genetic groups. Results for the different values of *K* were averaged using CLUMPAK’s [49] LargeKGreedy algorithm with 2 000 repeats. Selection of optimal number of *K* ancestral groups was verified with significant principal components estimated from scree plot using R package *LEA* v.3.6.0 [50].

For assignment and admixture estimates, 90% credible regions were produced along the runs for each individual. After deciding the most probable number of genetic clusters (*K*), an individual assignment probability (q) of ≥ 0.7 into a single cluster was set as a threshold of acceptance as ‘pure’. Thus, fish with lower than 0.7 assignment were considered as possible hybrids/admixed individuals between the supported genetic clusters.

Results from PCA and STRUCTURE clustering approaches were compared with each other and aligned. At this stage, suspected hybrids were removed and the supported *S. norvegicus* B species (see Results) combined. Then, the resulting species-level assignments were added to those 1990 individuals to enable PCA assignment for them. This was done because this dataset contained many fish from mixed catches and/or without any accompanying species assignment.

#### 2.7.3 Genetic differentiation between and within species

After all individuals were grouped into the species *S. mentella, S. viviparus, S. norvegicus* A and *S. norvegicus* B, the pairwise genetic differentiation between them (as *F*_ST_ defined by Weir & Cockerham [51]) was estimated using the *R* package *StAMPP* v.1.6.3 [52] with 1,999 bootstraps to calculate corresponding 95% confidence intervals and *p* values. Significance level was adjusted for multiple comparisons with FDR correction [53]. Pairwise matrices of differences were calculated as Nei’s genetic distance and visualized as dendrogram with *poppr,* v. 2.9.4 [54] with node supports obtained with 1 000 bootstraps.

Expected (*H_e_*) and observed (*H_o_*) heterozygosities together with inbreeding coefficients (*F*_IS_) were estimated with the *diveRsity* package, v. 1.9.90 [55]. Confidence intervals at 95% for *F*_IS_ was calculated with 1,000 bootstraps. Deviations from Hardy-Weinberg expectations (HWE) were estimated per population with *pegas* package, v. 1.3 [56] as the proportion of loci significantly deviating from HWE after MC permutation tests with 1 000 repeats and reference p-value FDR corrected for multiple testings.

#### 2.7.4 Investigating substructures within species

As higher levels of genetic structure can mask lower level of genetic hierarchies [48], the STRUCTURE run explained in chapter 2.7.2, was repeated also on species-level. Specifically, *S. mentella* and *S. norvegicus* B were analysed further, as they showed significant heterozygote deficiencies and large deviations from HWE (see Results), possibly suggesting population sub-structuring, i.e. Wahlund effect [57].

We estimated pairwise divergence as *F*_ST_ for within-species subgroups. Moreover, we defined the most separating loci using PC loadings in *factoextra* as explained above and checked locus independence with *R* package *poppr*, version 2.9.1 [54] by calculating the association indices over all loci in the dataset using 399 permutations, as well as per each locus pair. The latter was done ascertain that observed groupings did not arise from physical linkage between used markers (as observed e.g. in [58]).

#### 2.7.5 Estimation of assignment accuracy

Finally, we estimated how well selected SNP panels differentiated the species from each other using the assorted species data as baseline. We used *R* package *assignPOP* v 1.3.0 [59] to collect a subset containing 25%, 50%, or 75% of the used loci that had the highest differentiation (measured as *F*_ST_) between the baseline groups. In the training runs, a randomly selected 50, 70, or 90% of the individuals were included. Each resampling event was repeated 100 times. Assignment accuracy, i.e. fish correctly assigned into its original group, was then estimated via MC cross-validation procedure.

## 3. Results

### 3.1 Genome-wide divergence between species based on pool-seq data

The pool-seq data consisted of 357×10^6^ reads of *S. norvegicus* A, 379×10^6^ reads of *S. norvegicus* B, 352×10^6^ reads of *S. mentella* and 366×10^6^ reads of *S. viviparus*. Average insert size was 270bp and GC content 40.8%. Initial SNP calling returned a total of 7.4×10^6^ polymorphic sites across the reference genome of which 274,955 remained after quality and coverage filtering.

Based on this filtered genome-wide data, differentiation between pooled samples ranged from *F*_ST_ = 0.33 (*S. norvegicus* A vs *S. norvegicus* B) to *F*_ST_ = 0.61 (S. *norvegicus* A vs *S. viviparus*). Relative proximity between the two *norvegicus* samples was confirmed by PCA (Fig 2A). The fist principal axis separated *S. viviparus* sample from the others whereas the second axis separated *S. mentella*. The *F*_ST_ heatmap and associated dendrogram (Fig 2B) showed the same general pattern.

**Figure 2.**
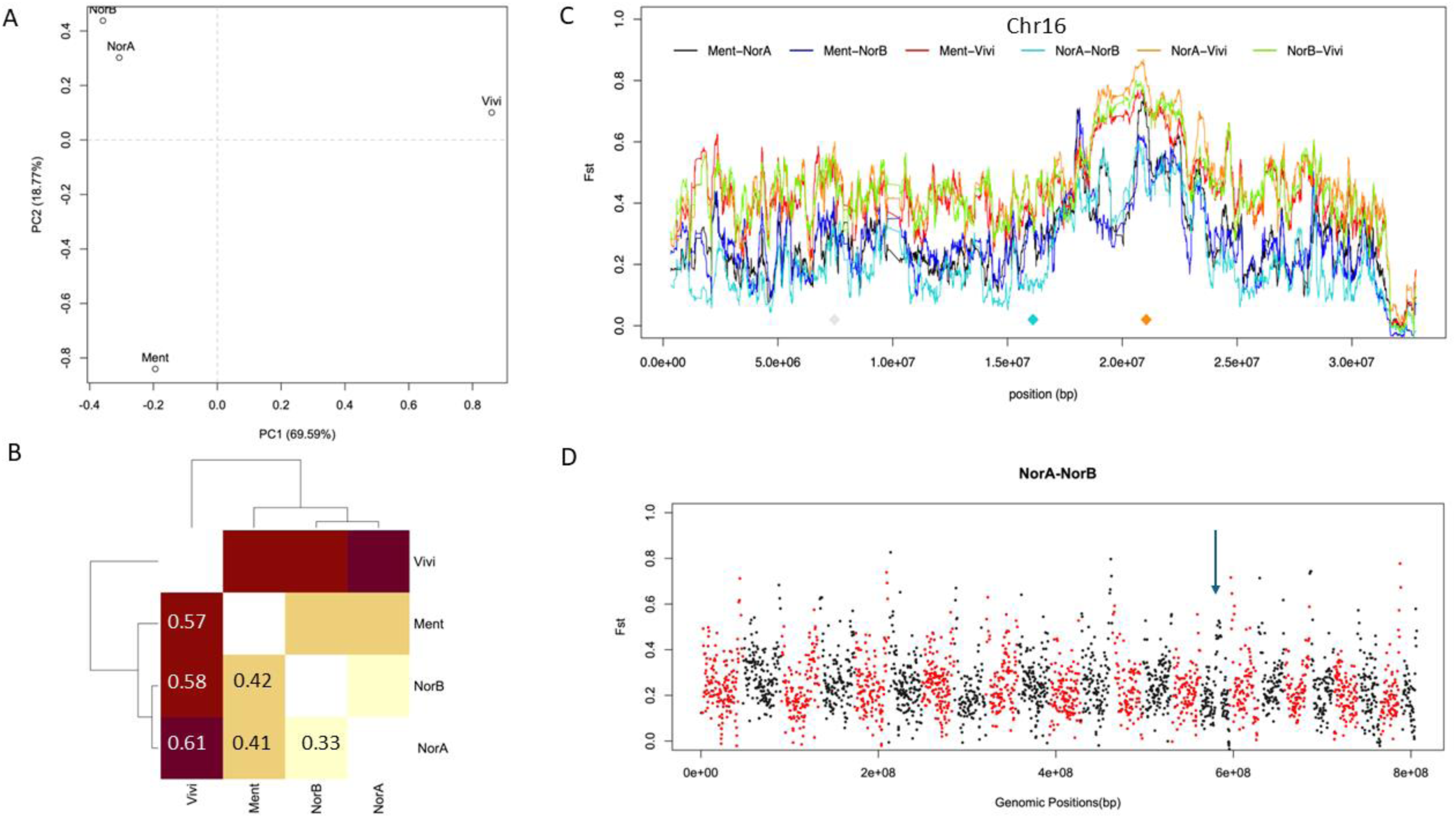
Genomic divergence between the northeast Atlantic *Sebastes* species based on pool-seq data. **A)** PCA for the species along the first two axes. PC1 explained 69.6% of the total variance, and PC2 18.8%. **B)** Mean pairwise differentiation as phylogeny heatmap with measured mean *F*_ST_ values indicated, **C)** An example figure of *F*_ST_ variation between species along a single chromosome (CM091696.1; named here as 16 due to its order in the reference) where different species pairs are colour-coded as shown in the legend above the figure. Coloured diamonds below correspond to positions of selected SNPs. Corresponding figures for all linkage groups are included as S. Figs 1. **D)** Genome-wide variation of divergence, where each dot corresponds to a measured *F*_ST_ for each SNP. Alternating colours indicate 24 assumed chromosomes in *S. fasciatus* reference genome against which our data was aligned. Arrow indicates position of chromosome 16 shown in panel C. Please note used abbreviations for the species: Vivi = *S. viviparus*, Ment = *S. mentella,* NorB = *S. norvegicus* B, NorA = *S. norvegicus* B.

Estimating genetic distance along the genome revealed high variability across genomic positions, with regions of very high (*F*_ST_ ≥0.80) and relatively low (*F*_ST_ ≤0.20) divergence as illustrated for example along chromosome 16 (linkage group CM091696; Fig 2C). Most high divergence regions were found from ends of linkage groups and observed patterns were generally similar between species pairs (S. Figs 1). Even the least diverged pair, the two cryptic *norvegicus* species, showed relatively high mean divergence and clear peaks of differentiation across almost all linkage groups (Fig. 2D).

### 3.2 Species identification and population structure based on SNP panel

Six duplicated genotypes and 20 fish with too much missing data were removed from the 56 SNPs dataset, whereas for the 41 SNPs dataset the corresponding numbers were 54 and 79, respectively. The final SNP datasets that passed all filtering steps and 56 SNPs had 1964 fish, whereas the dataset with 41 SNPs had a total of 2776 fish. 857 fish were thus exclusively analysed with the smaller set of markers. The final dataset of 56 loci had a missing data rate of 2.12%, and that of 41 SNPs 7.01%.

#### 3.2.1 Supported genetic groups and their geographic distribution based on clustering analysis

Depending on the clustering method used, between 4 and 5 major genetic groups were supported. Clustering based on principal components clearly separated four groups along the first two axes (Fig. 3A) that together explained 64.9% of the observed genetic variation. Comparing detected clusters with collected individual metadata (STable 2) indicated that major genetic clusters largely match morphological species identification (Fig 3). Most numerous was *Sebastes norvegicus* B (Table 2), which comprised over half of the samples (52.5%), followed by *S. mentella* (32.1%), *S. norvegicus* A (13.2%), and *S. viviparus* (2.2%). Four individuals appeared roughly in the middle of the supported species clusters in the PCA and were inspected further as possible hybrids, documented before for these species [60]. Larger dataset of 2776 fish genotyped with 41 SNPs showed consistent PCA pattern (S. Fig 2).

**Figure 3.**
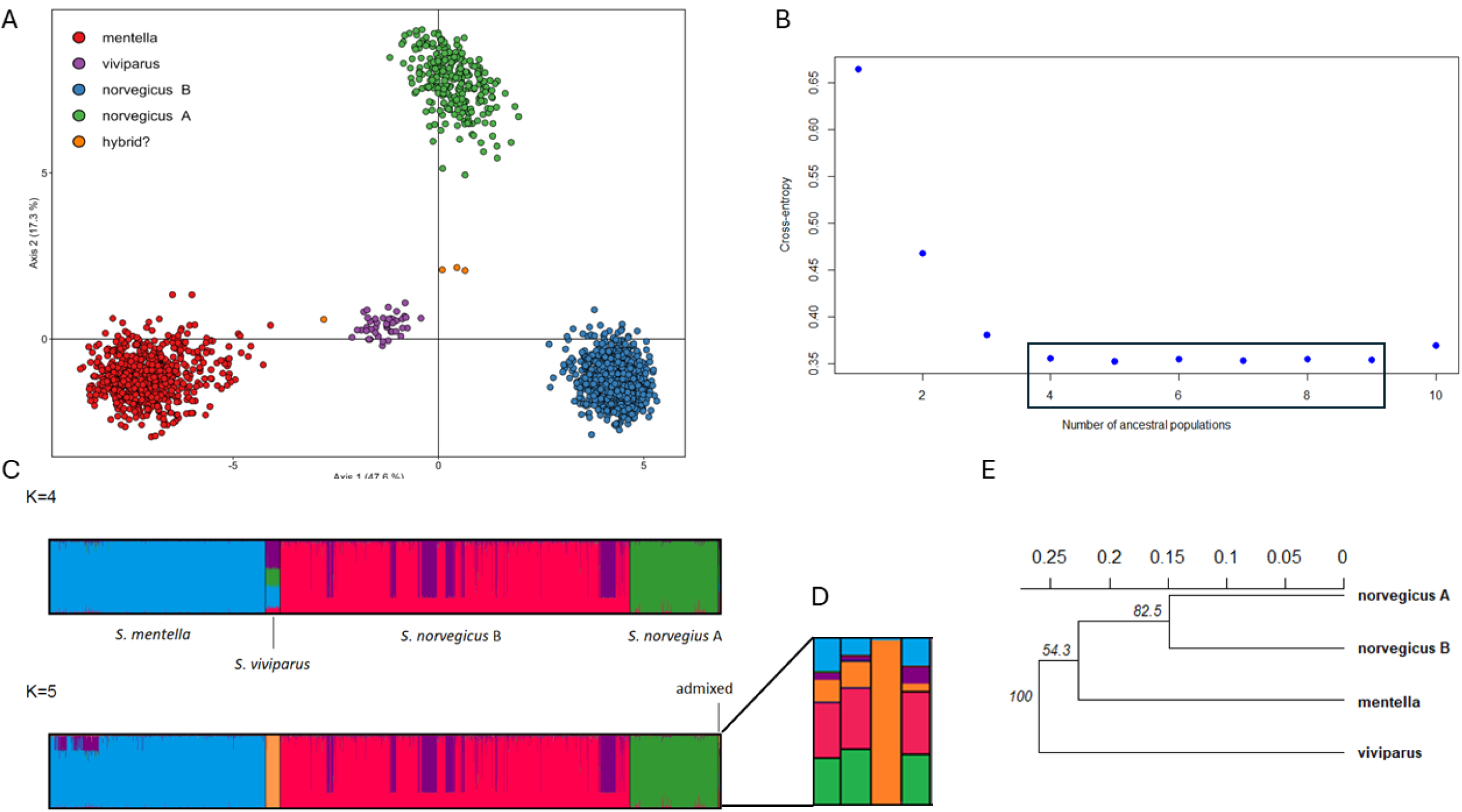
Major genetic groups of northeast Atlantic *Sebastes* species. Results are based on 56 selected SNPs and 1954 individuals. PCA and STRUCTURE clustering were conducted without any prior species information, which were added subsequently based on recorded field data. A) PCA results reveal species clusters and possible hybrids. B) Likely number of ancestral populations is 4-9 as supported by the lowest observed cross-entropy. C) The most likely number of genetic clusters is *K=*4-5 as suggested by Bayesian clustering approach. D) Zoom-in showing individual partitioning for those four fish suspected being hybrids. Note that colours for species between panels A and C/D are not consistent. E) Nei’s genetic distances as a dendrogram between the four species based on 1 000 bootstraps.

Some sub-structuring beyond species-level was evident in the datasets, as suggested by almost equally low values from 4 to 9 ancestral groups (Fig 3B), and STRUCTURE results of both datasets (Fig 3C, S. Figs 3-6). We deemed *K* = 5 as the most plausible number of genetic groups, as this was the highest number that created visually discrete clusters. It was also the best grouping according to Puechmaille statistics (S. Figs 4 and 6). Compared to PCA results, one additional group was supported within *S. norvegicus* B. This group appeared already at *K* = 4 (i.e. before all species appeared; Fig, 3C) supporting strong separation of this group. Additional levels of *K* were visually investigated and indicated within-species sub-structuring also in the *S. mentella* group. In contrast, the observed additional vertical group stratifications on roughly equal proportions (S. Fig. 7 and 10) is a typical artefact with no underlying population structure (see for example [61]).

**Figure 4.**
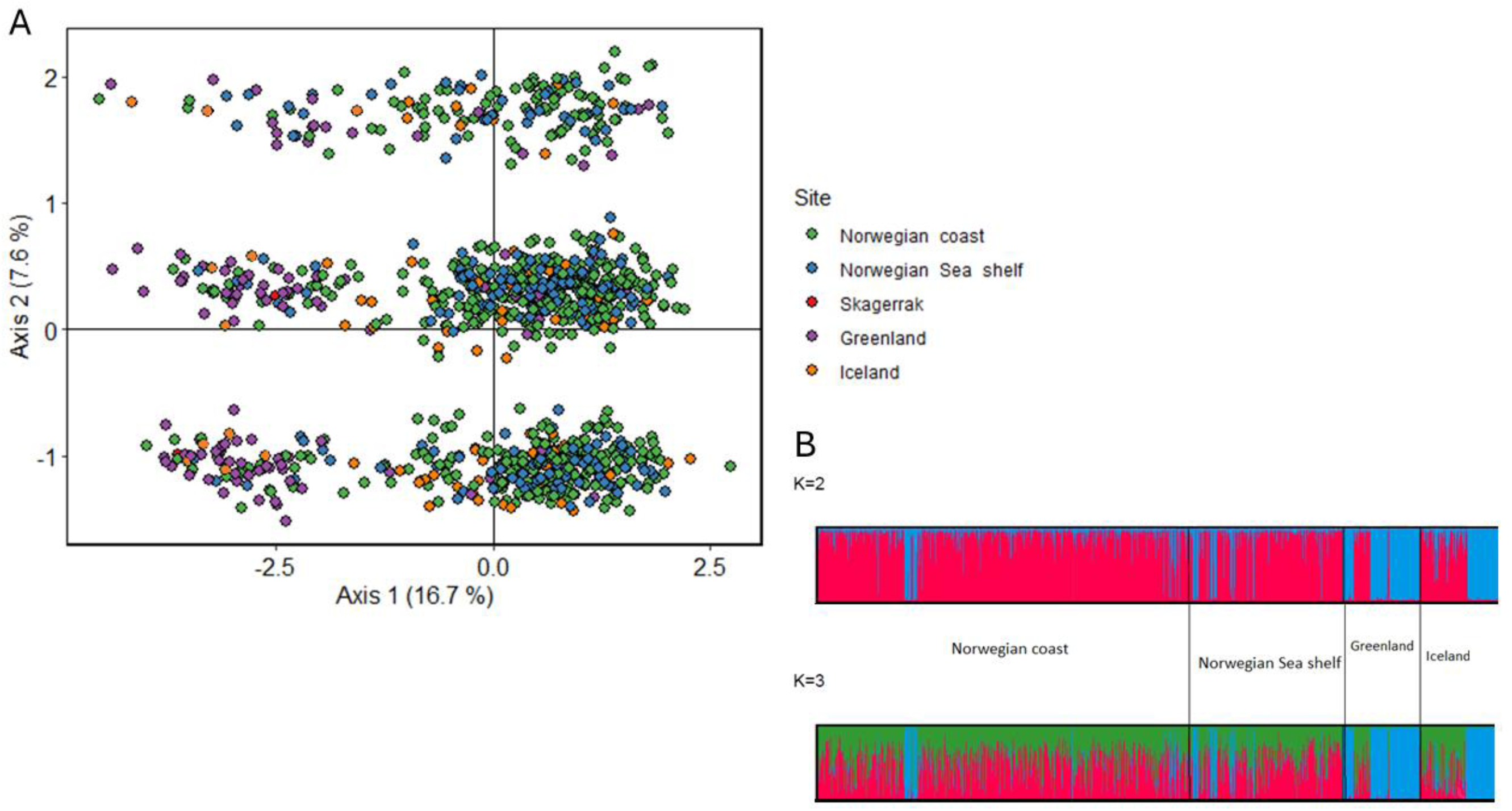
Genetic structure within *S. norvegicus B* based on 56 SNPs and 1033 individuals. **A)** PCA results showing two groups along axis 1 and a cryptic division along axis 2. **B)** STRUCTURE clustering results with clear separation between a mainly “Greenlandic” (blue) and mainly “Norwegian” (red) group.

Most species were found throughout our sampling range (STable 3; S. Fig. 7), but some spatial and temporal differences were apparent: Fish caught annually along the Norwegian coast as part of reference research cruises were comprised in large part of *S. norvegicus* B, whereas catches from Greenland, Iceland, and the Norwegian shelf were more mixed, as well as the small sample from Skagerrak. Samples from Lyngenfjord and the Norwegian Sea consisted (almost) entirely of *S. mentella*, whereas *S. viviparus* was least common overall but present in significant numbers (*N* ≥ 10) in samples from Greenland 2023, Norwegian shelf 2019, and Norwegian coast 2019. When interpreting distribution patterns of diverse *Sebastes* species in the northeast Atlantic, it is necessary to note that our sampling was not random, and thus these frequencies do not reflect the species’ abundancies.

#### 3.2.2 Genetic diversity and divergence among the species

Genetic diversity measured as expected heterozygosity (S. Table 4) was highest for *S. mentella* in both datasets (*H_obs_56loci_* = 0.202, *H*_obs_41loci_ = 0.171) and lowest for *S. viviparus* (*H_obs_56loci_* = 0.073, *H*_obs_41loci_ = 0.171). Heterozygosity deficiency was present in *S. mentella* (*F*_IS_ = 0.082, 95% CI: 0.047-0.166) and *S. norvegicus* B (*F*_IS_ = 0.098, 95% CI: 0.038-0.173) together with a relatively large proportion of loci not following expected Hardy-Weinberg proportions. Corresponding indices for the other two species, *S. norvegicus* A and *S. viviparus,* were not significant.

Pairwise genetic differences were high between all species, ranging from 0.633 to 0.757 when measured with 56 SNPs and from 0.714 to 0.789 with 41 SNPs (STable 4). Divergence was highly significant (*p* ≈ 0) in all cases. A dendrogram based on Nei’s genetic distance indicated shortest distance between *S. norvegicu*s A and B, connecting further to *S. mentella* and *S. viviparus* being the most distant (Fig. 1E). Node support was high (82.5%) for the separation of the *S. norvegicus* group, whereas the more basal split of *S. mentella* was less supported.

#### 3.2.3 Supported substructures and detected potential hybrids

Besides the four known species, further sub-structuring was supported for two species, *S. norvegicu*s B and *S. mentella* as indicated already indirectly by the deficiency of heterozygotes (see above), and directly by additional divisions supported by STRUCTURE analysis and phylogeny splitting (Fig. 3C, S. Figs 7-11). Therefore, these species were investigated in detail separately, and the obtained results are presented below.

##### *S. norvegicus* B

Among the 1033 and 1742 fish (56 and 41 SNPs datasets, respectively) assigned to *S. norvegicus* B, strongly supported subdivision into two groups was found. Individuals were mainly divided geographically along the first axis (Fig. 4A), where “western” type individuals were found mainly outside Greenlandic waters and Reykjanes ridge in Iceland (blue bars in Fig. 4B; S. Fig. 8), but also to lesser extent also along the Norwegian coast and the Norwegian Sea shelf (but with frequency of < 20%). In total, 183 fish belonged to this group with an individual assignment (*q*) of ≥ 0.7; 786 fish were assigned to the other group, and 58 were admixed/uncertain (for results with 41 SNPs data and 1742 fish, please see S. Fig. 7). When we compared this grouping with the known reference data, this western type was confirmed as “giants” *S. norvegicus*. Along the Norwegian coast, these giants were mainly documented furthest offshore compared with the more common type (S. Fig. 8). The constructed UPGMA tree based on Nei’s genetic differences also strongly supported this subdivision (S. Fig. 9). Measured pairwise *F*_ST_ between the two groups was 0.213 (*p ≈* 0). Genetic diversity, *H_exp_*, was 0.082 in “giant” cluster (*H_obs_* = 0.079, *F*_IS_ = 0.029 (*NS*)), and 0.105 in the more common type (*H_obs_* = 0.103, *F*_IS_ = 0.065 (*NS*)).

There was evidence of possible further cryptic substructure, as indicated by further stratification of PCA along the second axis (Fig. 4A) and uneven vertical subdivision with *K* = 3 (Fig. 4B). Such pattern could be connected to structural variation and physically linked SNPs (see e.g. [58]). We excluded this possibility, however, as the loci were originally selected from different contigs (see section 2.3), and did not show significant linkage (S. Figs. 11). Thus, we deem two genetic groups within *S. norvegicus* B as the predominant structure but cannot rule out an additional genetic hierarchy.

##### S. mentella

634 fish were assigned to *S. mentella* with 56 SNPs, and an additional 21 when 41 SNPs were considered. In a similar manner to *S. norvegicus* B, two to three genetic groups were supported (Fig. 5, S. Fig. 10-11), one of which was predominant in Greenland (62.5% of *S. mentella* there) but found sporadically elsewhere as well. Notably, all three *S. mentella* caught in Skagerrak were of this “Greenlandic” type (cluster 1). In total, 99 fish were assigned to this group, 516 to the other group (cluster 2; pink bars in Fig. 5B with *K* = 2), whereas 16 fish could not be assigned unambiguously to either, as they showed admixed genotypes. *F*_ST_ between the two distinct genetic groups was 0.148 (*p ≈* 0), and the split was highly supported in phylogeny (S. Fig 7). Genetic diversity measured as *H_exp_* was 0.204 in the “Greenlandic” cluster (*H_obs_* = 0.202, *F*_IS_ = 0.051 (*NS*)), and 0.190 in the other group (*H_obs_* = 0.178, *F*_IS_ = 0.048 (*\**)).

**Figure 5.**
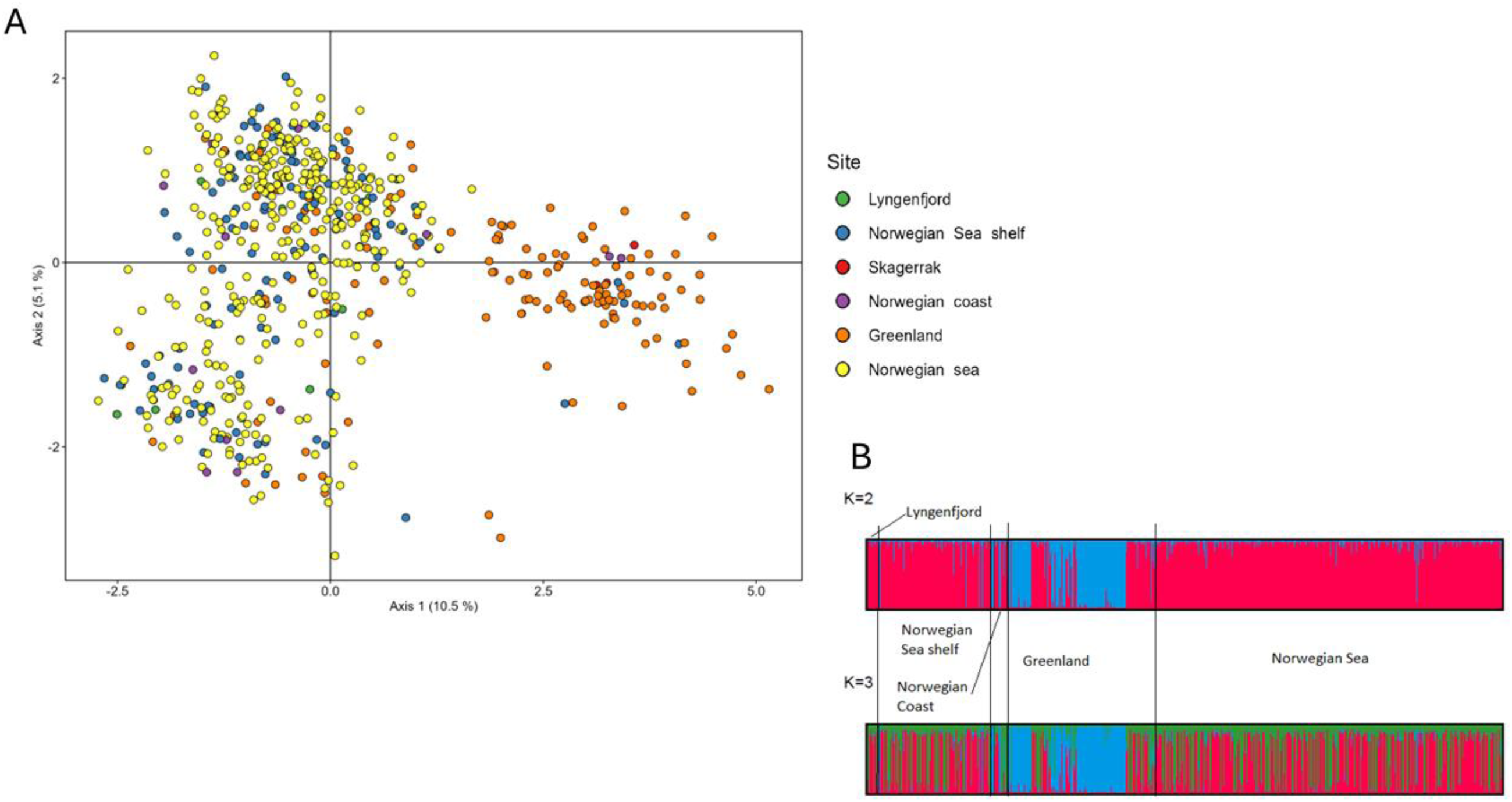
Genetic structure within *S. mentella* based on 56 SNPs and 634 individuals. **A)** PCA showing a strong, geographic separation along axis 1 and a more subtle separation along axis 2. **B)** STRUCTURE clustering analysis resolving similar groups.

When three groups were considered (*K* = 3) instead of two, an obscure pattern appeared among fish not belonging to the ‘Greenlandic’ cluster (Fig. 5B). This division is also apparent in the PCA as a diagonal split (along axis 2 in the cluster on the left side). This cryptic division (seen as green bars in Fig. 5B) received equal support to *K* = 2 in both datasets (S. Fig. 9) and was not due to linkage between used loci (S. Fig 8b). Also, the second cluster showed significant heterozygote deficiency (see above) further supporting substructure. *F*_ST_ between this cryptic division was 0.065 but not significant (*p* = 0.353). Thus, while it is likely that *S. mentella* consists of three genetic entities (as previously found by [18]), results of this dataset are inconclusive. Fifteen loci were specifically selected to investigate the population structure within *S. mentella*, and when only they were selected, and *F*_ST_ recalculated, the divergence was clearly higher between all clusters but yet again not significant between the cryptic clusters 2 and 3 (*F*_ST_1vs2_ = 0.254, *p ≈* 0, *F*_ST_1vs3_= 0.146, *p ≈* 0, *F*_ST_2vs3_= 0.153, *p* = 0.247).

##### Hybrids

Four fish had intermediate genotypes that placed them outside well-defined clusters (i.e. species) in the PCA (Fig. 3A). Three of them (0.1%) were identified as possible hybrids (Fig. 3D) because of their highly admixed genotypes. Two were caught in the Norwegian coast (in 2020 and 2022), and one on the Norwegian shelf (in 2016).

#### 3.2.4 Assignment accuracy of the SNP panels

The selected loci had high assignment accuracy on the species level (S. Figs 12). When at least half of the loci were (randomly) included, all fish were correctly assigned. Only when 25% or below of loci (*N*≤14) were included, some erroneous assignments were observed. We tested assignment power also on a lower genetic structure level by including subdivision of *S. norvegicus* B. While this still resolved the species well, mean accuracy for the subdivision among *S. norvegicus* dropped somewhat, and many individuals originally assigned into the ‘giant’ (*S. norvegicus* B subgroup) cluster were assigned incorrectly.

## 4. Discussion

Here, we investigated genetic divergence among and within the northeast Atlantic *Sebastes* species using two approaches. The first method, pooled sequencing of individuals is a cost-effective approach to compare full genomes [8, 29] and source markers for further use [24]. It also allows to interrogate the genomic landscape of divergence, which may help understand the mechanisms of speciation [62, 63] and other types of genetic differentiation [64]. We utilized a recently published, high-quality reference genome from a closely related [9] northwestern Atlantic species, *S. fasciatus,* to align our sequences against. Our comparison of genome-wide differences (*F*_ST_) revealed high mean divergence between all species pairs - including the cryptic pair *S. norvegicus* A and B - and thus confirms their genetic distinctiveness. Compared with previous microsatellite-based estimates of divergence [13], the estimates obtained here were two to four-fold higher. This could be due to many factors: e.g. stringent filtering criteria was adopted [65], the reference genome used was from another species [66], and the fact that these markers are inherently different [67]. Nevertheless, we ascertain large divergence between all North Atlantic *Sebastes* species and across their entire genomes. Another issue worth mentioning in the context is the significance of reference genome improvement: The SNP panels were developed using the same pool-seq data as in this study, but with much more fragmented reference genome with over 75k scaffolds (GCA_900302655.1; for details see [24]), and the sourced discriminatory SNPs seem to mostly represent local peaks of divergence but rarely hit the most diverged parts of each chromosome (S. Figs 1).

Interestingly, variation patterns along chromosomes appear highly synchronous among species (S. Figs 1), with only magnitude varying depending on species pair. This implies that the genome organization itself likely largely defines the general differentiation patterns. Variation in recombination rate and gene positions could, e.g. be such determining factors [62, 68]. Regions of clearly elevated differentiation were mostly observed at the ends of chromosomes, possibly due to their association with e.g. telomeric and/or pericentromeric chromatin regions [69]. An early karyotype study of related species from the *Sebastes* genus [70] suggested conserved genomic structure with all but one chromosome being acrocentric, also fitting with the idea of divergence accumulation due to structural determinants [71]. However, as we currently lack high-quality reference genomes together with information of these species’ chromosomal karyotypes, recombination rates, and gene locations, this remains speculative for now.

The SNP panel used here proved highly discriminatory - despite missing most peaks of divergence - and suitable to detect officially acknowledged species, but also the cryptic *S. norvegicus* species pair A and B, and the giant morph which based on our SNP analyses here seems to be closely related to *S. norvegicus* B (S. Fig. 9). The distinction of fast-growing, large morph was suggested already in the 1960s [27] and has later been supported by morphological and molecular studies [13, 72]. Among the samples analysed here, the giant type was mainly detected in Greenland, but also in Reykjanes Ridge in Iceland (from where it was described for the first time). Also, for the first time ever, the giants were found on the Norwegian coast and continental shelf. Interestingly, both *S. norvegicus* individuals caught in Skagerrak were also confirmed to be giants (S. Table 3). Compared with the other *S. norvegicus* B type, i.e. the one that is commonly considered to be golden redfish, giants were collected mainly further offshore (S. Fig. 8), along the continental shelf break and in the Barents Sea, suggesting possibly at least partial spatial separation between these two types. The cryptic species, *S. norvegicus* A, in contrast, had largely overlapping distribution ranges with golden redfish (i.e. non-giant *S. norvegicus* B in our analysis) and *S. viviparus* as well (S. Fig. 7) suggesting that other mechanisms besides simple spatial differentiation are likely keeping these species apart. A recent morphometric study by Bruvold *et al*. (2025) showed that *S. norvegicus* A and B are phenotypically also diverged: especially beak length, eye diameter, pectoral fin length, and caudal peduncle height were significantly different between the two. Consistently with observations by Bruvold *et al*. (2025), *S. norvegicus* A was clearly less prevalent (∼10%) among our samples compared to *S. norvegicus* B (>50%). Whether this reflects true population size or distribution differences, or for example seasonally variable catchability probabilities (and thus also possible temporal and/or spatial separation), remains to be studied. Considering that redfish are known for their wide ecological variability [9], several options behind this genetic separation are plausible [11]. For example, any differences in preferences related to breeding ecology could easily lead to separate gene pools and divergence even in sympatry [73, 74]. Such ecological speciation processes may co-occur with genomic changes that can uphold this separation [73, 75].

We detected genetic sub-structuring also within *S. mentella*, concordant with previously documented depth-related ecotypes [11, 14, 18, 25]. Our samples were mainly split along the east-west axis (Fig.5, S. Fig. 11) with vast majority of individuals from Greenland belonging into one group, assumed ‘slope’ ecotype (as presented in Saha *et al.* 2021), whereas fish caught on the east side of the Atlantic mainly formed the other, assumed ‘pelagic’ ecotype (Figs 5). This genetic split was strongly supported (S. Fig. 9). Within this northeastern Atlantic group, mainly consisting of fish caught from deep waters in the Norwegian shelf and Norwegian Sea, additional, more subtle subdivision was observed (Fig. 5A) which may represent the division between ‘shallow pelagic’ and ‘deep pelagic’ types. Additional splitting was reported also by Saha *et al.* (2021) attributed largely differentiation between Irminger Sea vs the Faroe Islands. However, these areas were not included in our study, and we could not associate detected differentiation beyond the assumed pelagic and slope groups. Accordingly, it could be possible that ‘pelagic’ group split is indeed driven by some other factor such as e.g. depth. Interestingly, Benestan and colleagues [11] also reported unidentified genetic grouping among *S. mentella* samples on the northwestern Atlantic. Notably, these groups showed spatial overlap in this study (Fig. 5) indicating that geographic location alone is not the decisive dividing factor, but rather that these groups mix in space.

Hybridization between *Sebastes* genus species has repeatedly been documented in the North Atlantic [17, 76, 77] and elsewhere [78, 79], indicating that species barriers within this genus are not definite. The reported North Atlantic cases are hybrids between *S. mentella* and other species, mainly *S. fasciatus* which was not included in this study. An interesting perspective related to hybridization and introgression was added by Benestan *et al.* (2021) who used genome-wide markers to investigate West Atlantic *Sebastes* species, *S. mentella* and *S. fasciatus*, and did not find any signs of recent hybridization between them. The authors suggest that introgression events are likely ancient and therefore not limited the establishment of barriers to gene flow between the species. We detected three individuals among our samples (∼0.1% of all) that were not genetically ‘pure’ representants of any *Sebastes* species or ecotypes found and described here but instead seemed highly admixed (Fig 3D). Observed admixture patterns did not support F1 hybridisation patterns either (i.e. with about 50:50 of two types, i.e. colours in the STRUCTURE bar plot) as these fish seemed to have genetic admixture from all species. This is somewhat unexpected considering these markers are highly discriminatory between these species (S. Table 4). There are a few possible explanations: First, these fish could be some sort of back crosses, possibly even involving more than two species. However, this would be more plausible in situations where barriers of gene flow are weak and hybrids occur in great numbers (like shown for *S. fasciatus* and *S. mentella* in the Canadian waters [17] and for two Pacific *Sebastes* species [79]) which is not the case here. Alternatively, the admixed fish could represent another distinct genetic entity: Bayesian clustering methods tend to divide individuals into existing genetic groups somewhat evenly if the true group is not accounted for (and thus minimizing deviation from the model assumptions; [80]). This can be seen from our species-level grouping as well (Fig. 3C): when too few clusters were included in the model (i.e. *K* was assumed to be 4 and not 5), all *S. viviparus* appeared highly admixed while they are in fact a highly distinct species. If this is the case here and there is indeed one more genetic group present, albeit in very low numbers, it would suggest that North Atlantic *Sebastes* species do not likely hybridize. Future studies involving full genomes will shed light on these, yet unanswered questions of *Sebastes* species differentiation in the North Atlantic. But when it comes to species monitoring and management - as few as two to three SNPs are enough to identify species reliably [11, 24]. This will come in handy especially for further investigations of the cryptic *S. norvegicus* species which are currently sourced and managed as a single stock.

## Supporting information

Jansson_etal_2025_Sebastes_supplements

## Acknowledgements

We express our gratitude to personnel onboard during the IMR surveys together with commercial fishers for collecting all samples used in this study. A special thanks to Martin Molander and colleagues for collecting our first *Sebastes* samples from Skagerrak. We thank Matthew Kent, CIGENE, and NorSeq Sequencing core (Ullevål) & Functional Genomics Group Department of Medical Genetics at the Oslo University Hospital for providing the sequences for this study. Warm thanks also to Laila Unneland for her assistance in the laboratory.

## Author contributions

TJ acquired funding for this study, gave the original idea for the study, coauthored the paper and is the corresponding author

EJ wrote the original draft and was responsible of SNP analyses and most visualizations included

FB was responsible for pool-seq data handling, analysis and illustrations and coauthored the paper

HC had advisory role, provided samples from Greenland and coauthored the paper AS had advisory role and coauthored this paper

CAT was responsible for map illustrations and data organization together with co-authoring this paper

IMB had advisory role and coauthored this paper

AMR was responsible for SNP genotyping and coauthored the paper

## Data availability

Data on which this study is based on will be published upon final acceptance.

