## Supplementary material for "Genome-wide differentiation and SNP-based identification of northeastern Atlantic *Sebastes* species": Jansson_etal_2025_Sebastes_supplements

**Supplementary data for study " Genome-wide differentiation and SNP-based identification of northeastern Atlantic *Sebastes* species" by Jansson *et al.* (2025)**

**Supplementary Figures 1. Genomic landscape of divergence ($F_{ST}$)** between northeastern Atlantic *Sebastes* species aligned against the used reference genome (GCA_043250625.1; https://www.ncbi.nlm.nih.gov/datasets/genome/GCA_043250625.1/). Assumed chromosomes are shown below in the order they appear in the reference. Pairwise $F_{ST}$ was measured in sliding windows of 100 SNPs. Different comparisons are shown with different colours as indicated above each figure. Diamonds on bottom of figures show genomic positions of SNPs in used panels in this study. Their colour indicates for which species-pair comparison each SNP was specifically selected for.

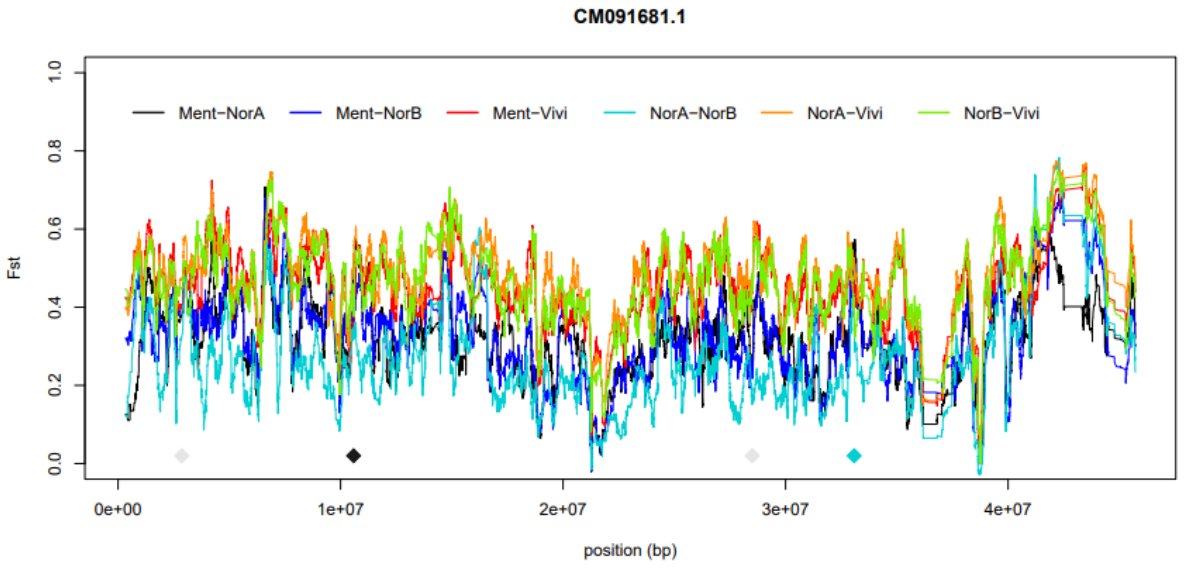


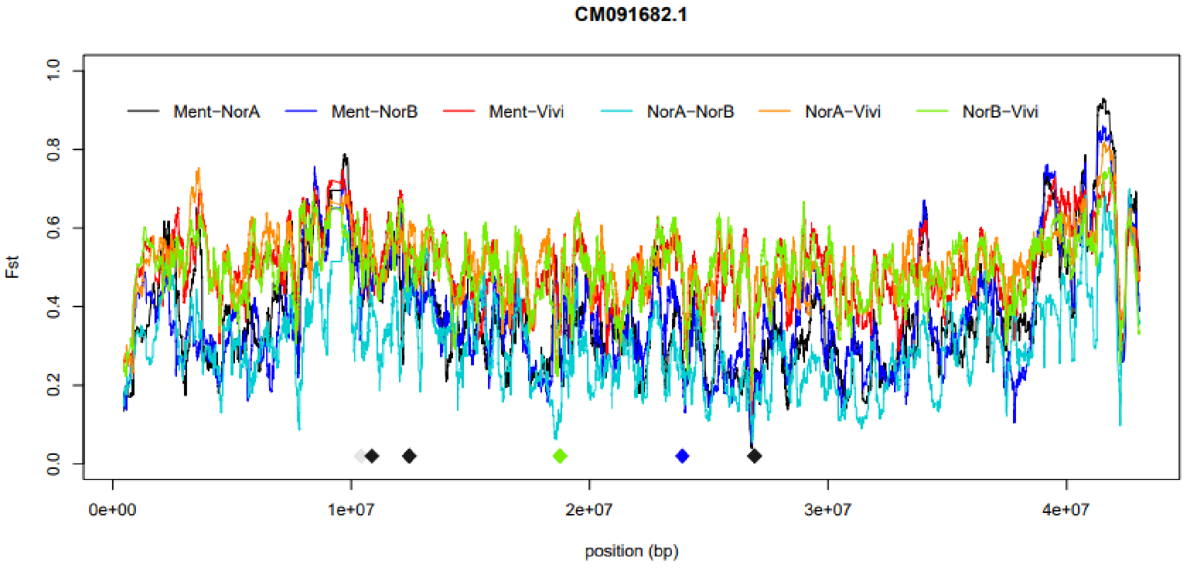

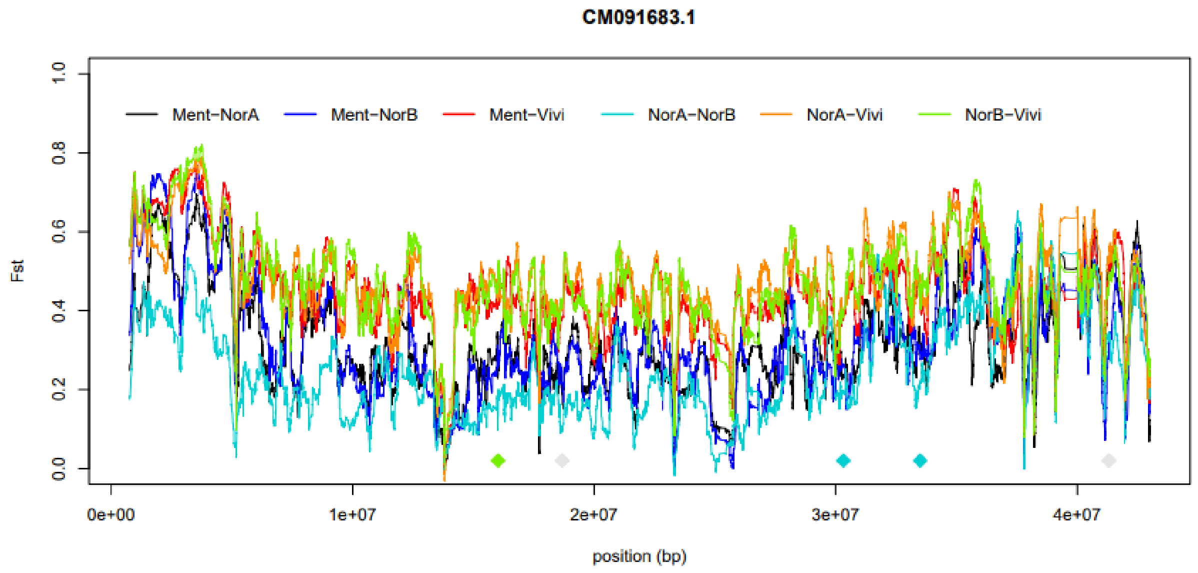

CM091683.1
Ment−NorA
Ment−NorB
Ment−Vivi
NorA−NorB
NorA−Vivi
NorB−Vivi
Fst
0.0
0.2
0.4
0.6
0.8
1.0
0e+00
1e+07
2e+07
3e+07
4e+07
position (bp)


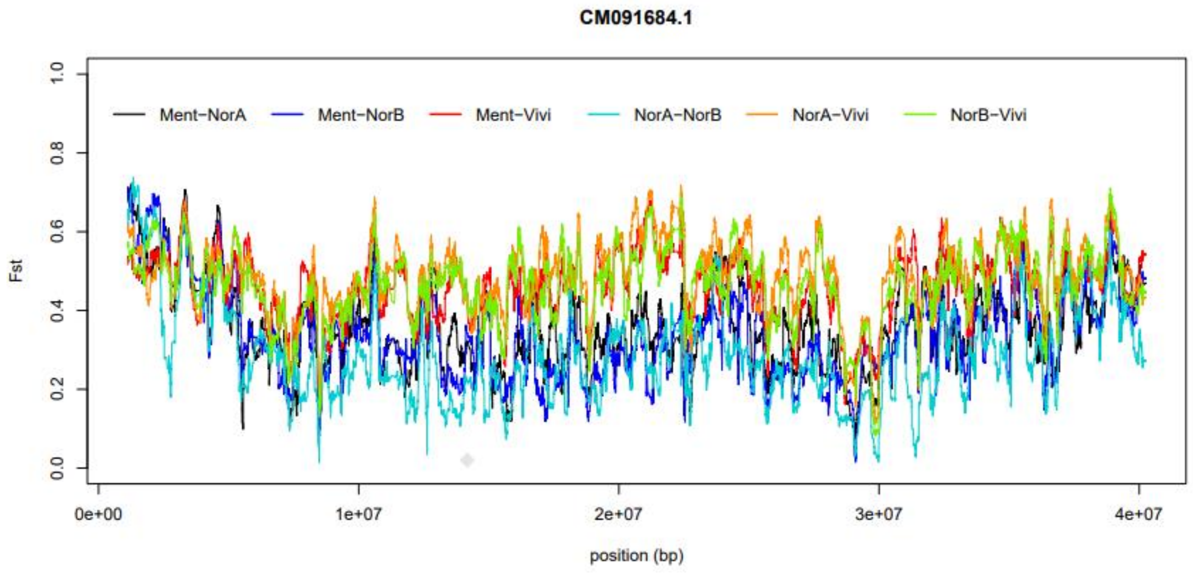

CM091684.1
Ment−NorA
Ment−NorB
Ment−Vivi
NorA−NorB
NorA−Vivi
NorB−Vivi
Fst
0.0
0.2
0.4
0.6
0.8
1.0
0e+00
1e+07
2e+07
3e+07
4e+07
position (bp)


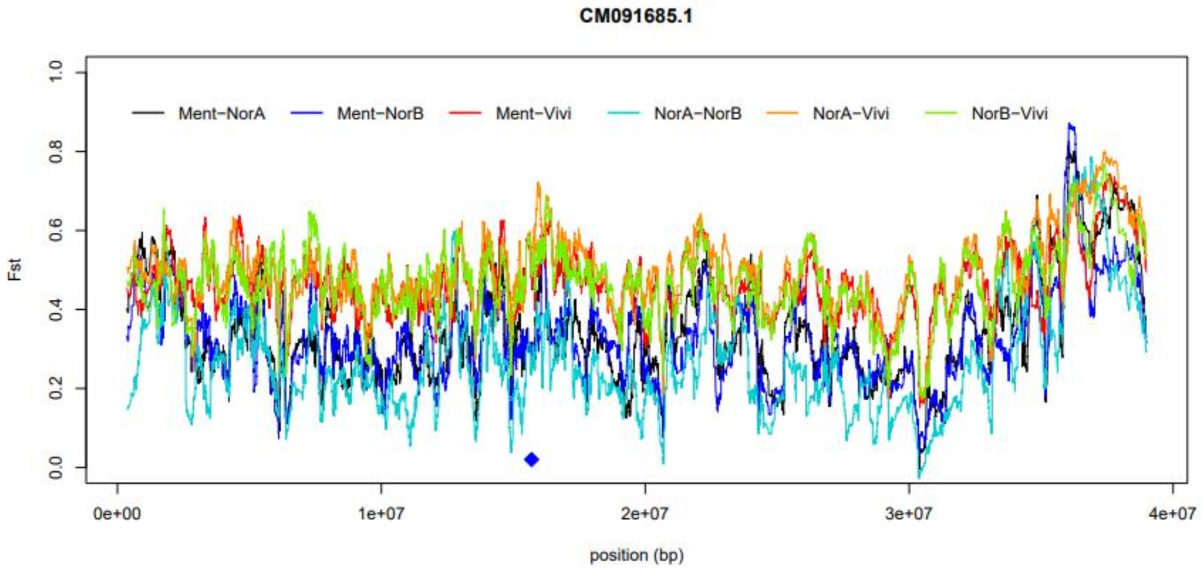

CM091685.1
Ment−NorA
Ment−NorB
Ment−Vivi
NorA−NorB
NorA−Vivi
NorB−Vivi
Fst
0.0
0.2
0.4
0.6
0.8
1.0
0e+00
1e+07
2e+07
3e+07
4e+07
position (bp)

**CM091686.1**

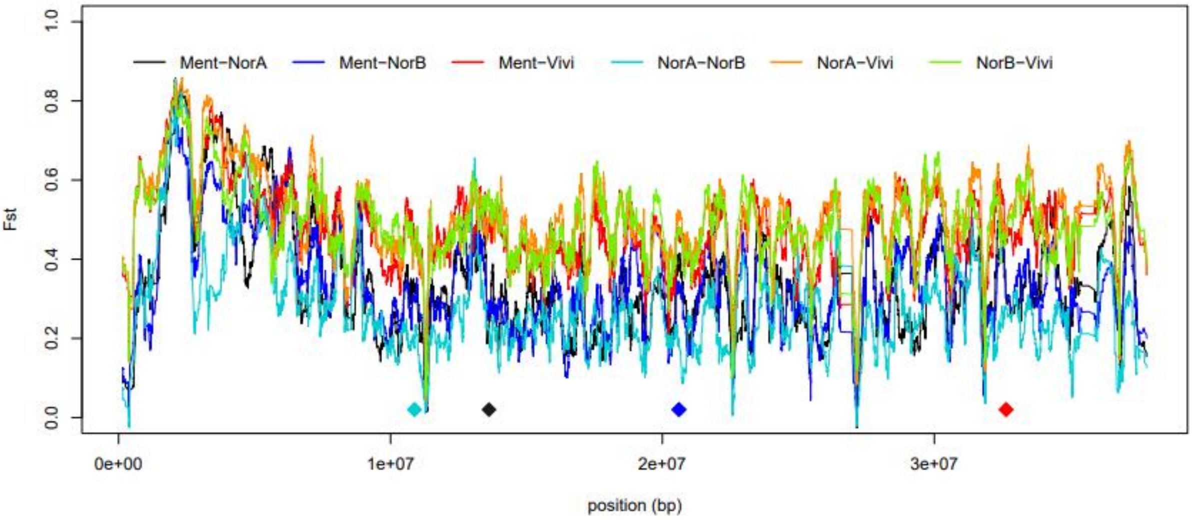


**CM091687.1**

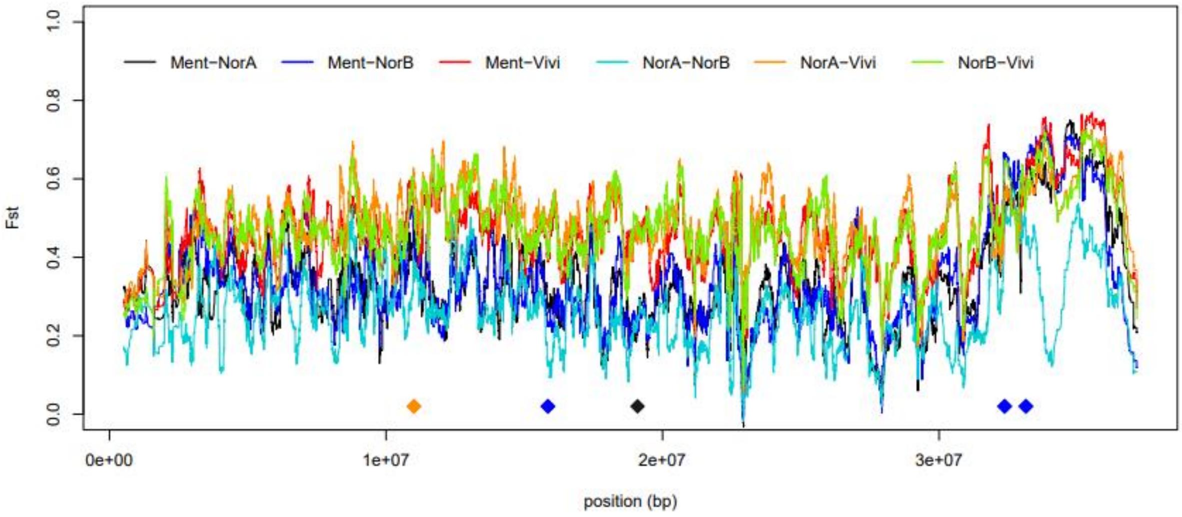


**CM091688.1**

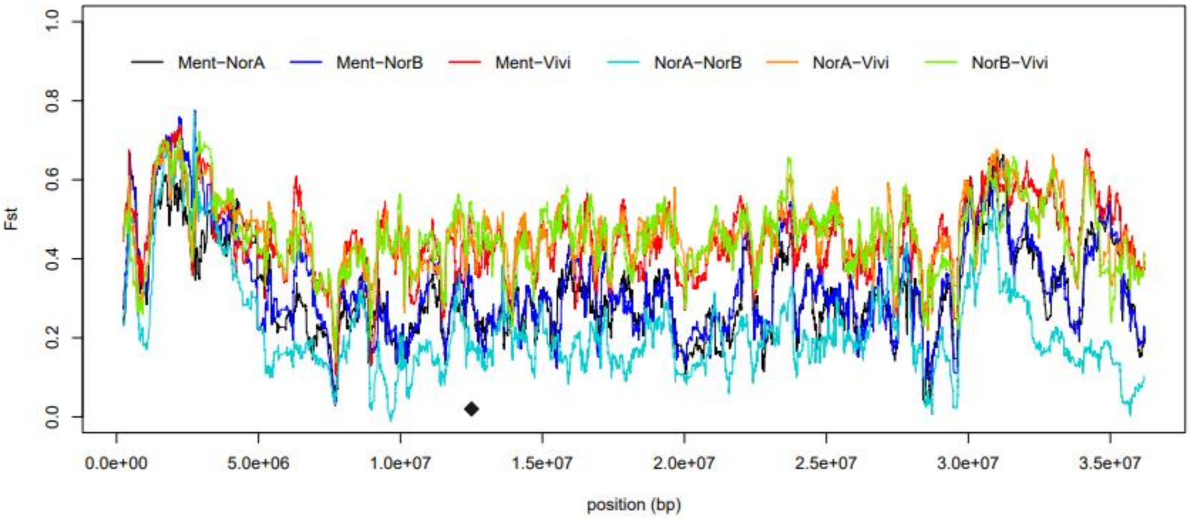

CM091689.1

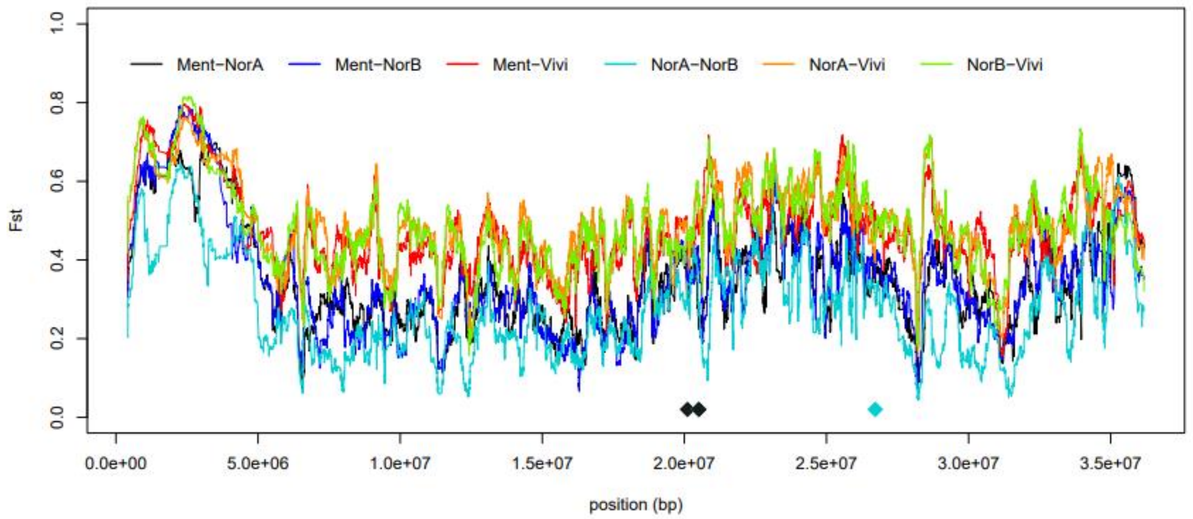


CM091690.1

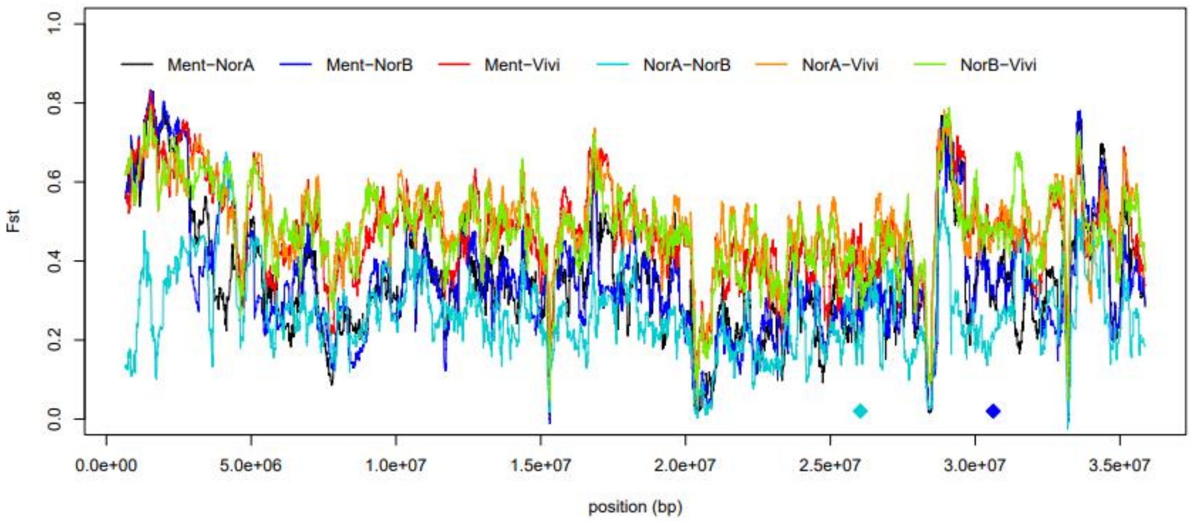


CM091691.1

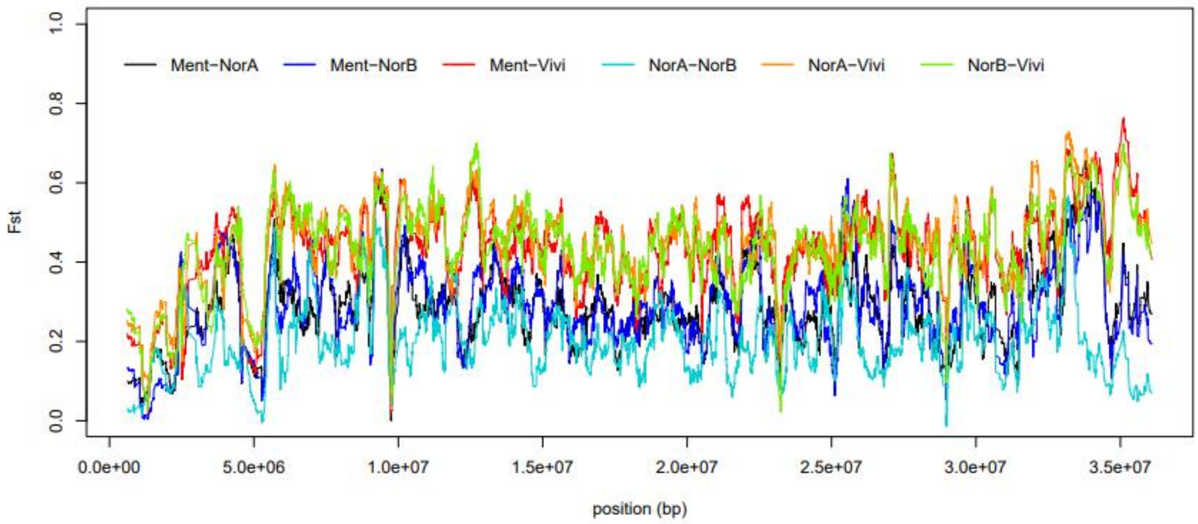

**CM091692.1**

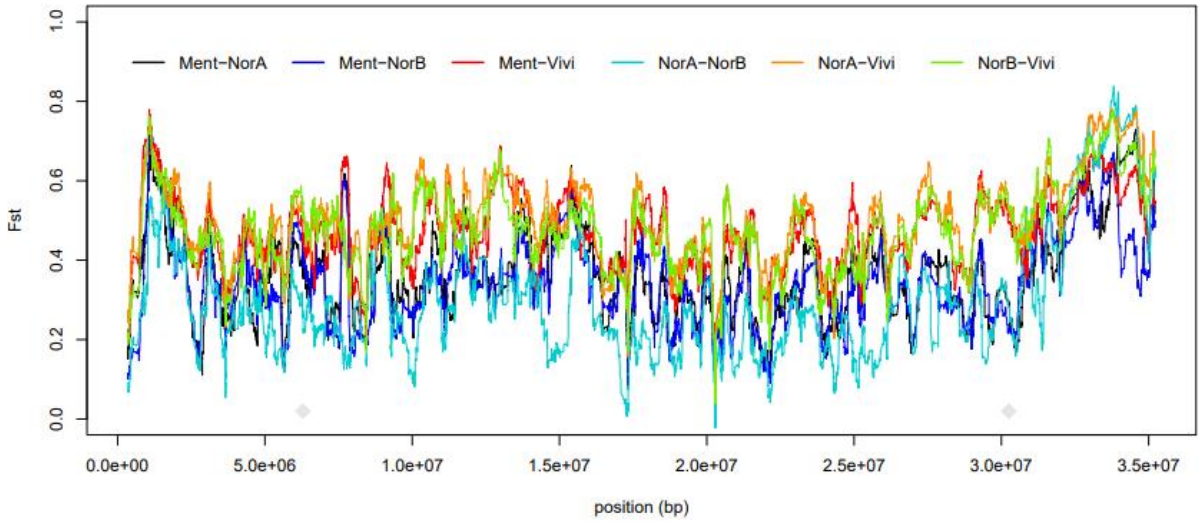


**CM091693.1**

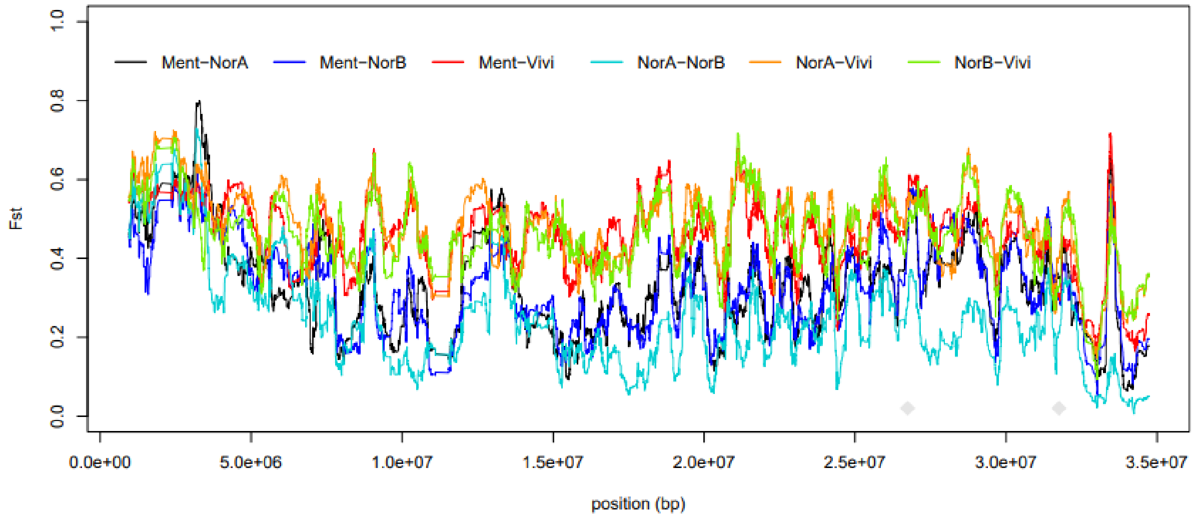


**CM091694.1**

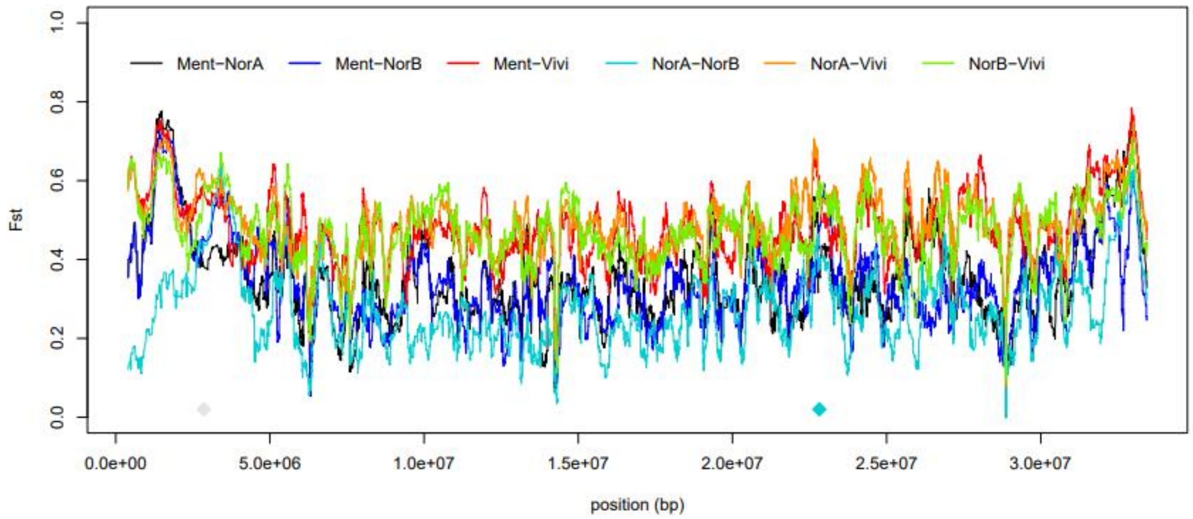

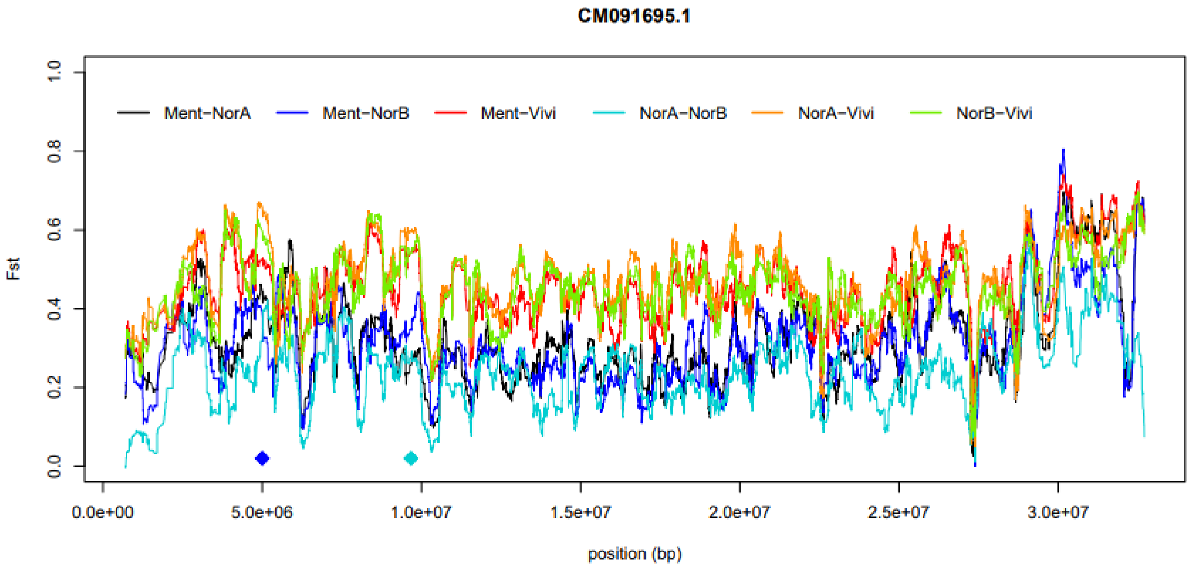
CM091695.1
Ment−NorA
Ment−NorB
Ment−Vivi
NorA−NorB
NorA−Vivi
NorB−Vivi
Fst
0.0
0.2
0.4
0.6
0.8
1.0
0.0e+00
5.0e+06
1.0e+07
1.5e+07
2.0e+07
2.5e+07
3.0e+07
position (bp)

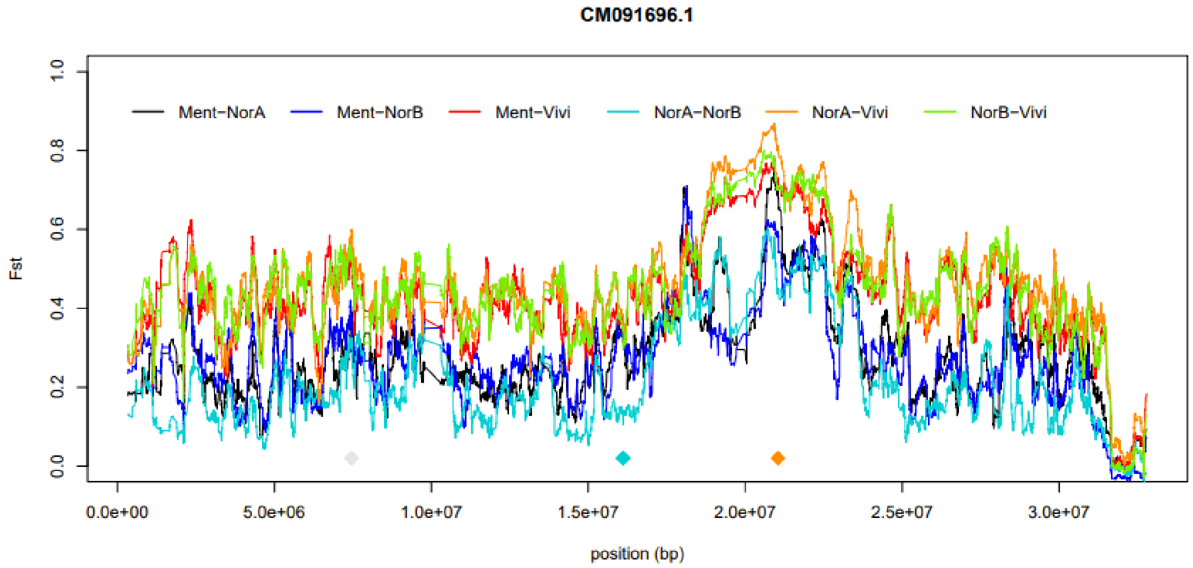
CM091696.1
Ment−NorA
Ment−NorB
Ment−Vivi
NorA−NorB
NorA−Vivi
NorB−Vivi
Fst
0.0
0.2
0.4
0.6
0.8
1.0
0.0e+00
5.0e+06
1.0e+07
1.5e+07
2.0e+07
2.5e+07
3.0e+07
position (bp)

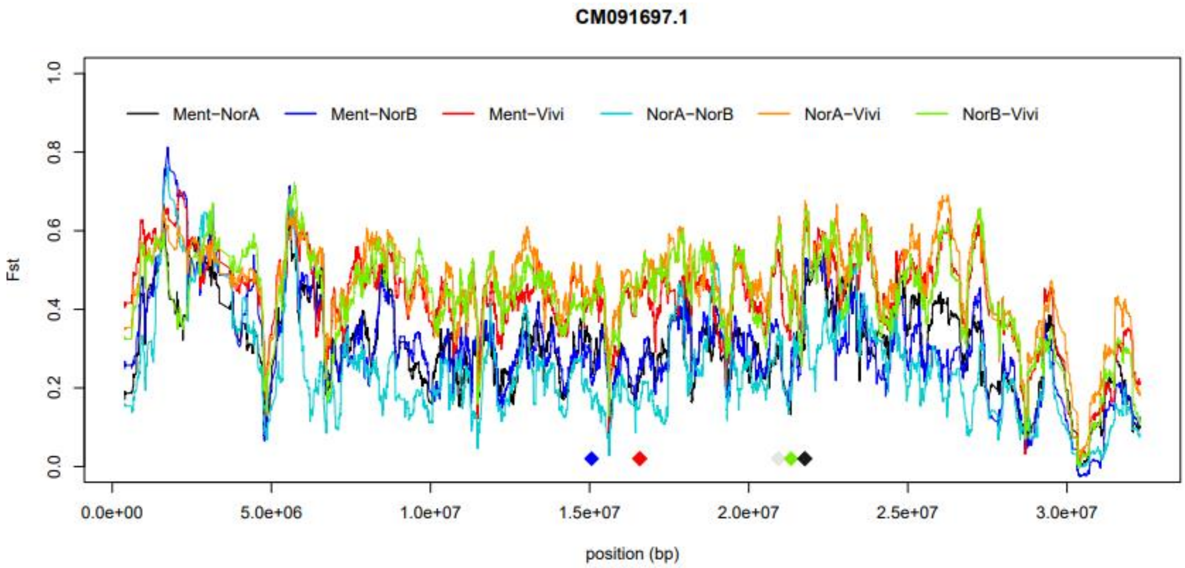
CM091697.1
Ment−NorA
Ment−NorB
Ment−Vivi
NorA−NorB
NorA−Vivi
NorB−Vivi
Fst
0.0
0.2
0.4
0.6
0.8
1.0
0.0e+00
5.0e+06
1.0e+07
1.5e+07
2.0e+07
2.5e+07
3.0e+07
position (bp)

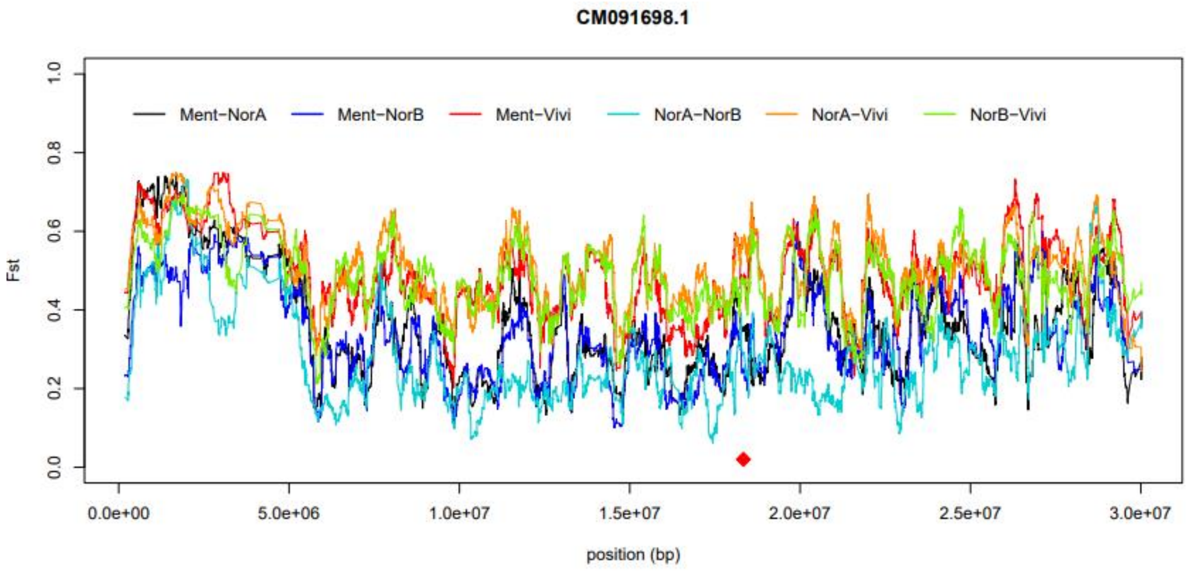
CM091698.1
Ment−NorA
Ment−NorB
Ment−Vivi
NorA−NorB
NorA−Vivi
NorB−Vivi
Fst
0.0
0.2
0.4
0.6
0.8
1.0
0.0e+00
5.0e+06
1.0e+07
1.5e+07
2.0e+07
2.5e+07
3.0e+07
position (bp)

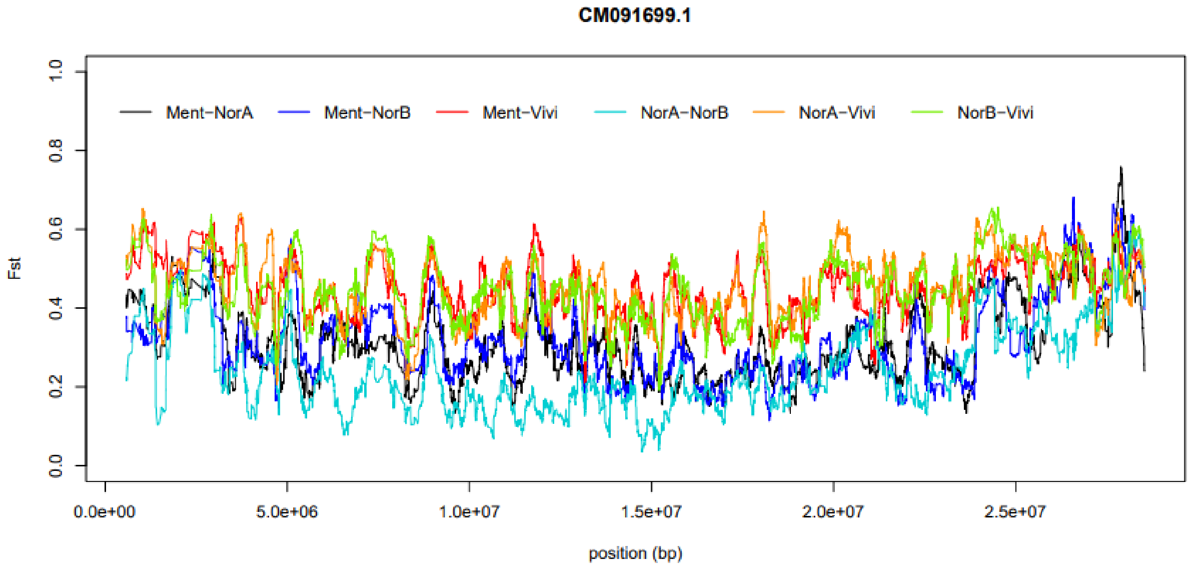
CM091699.1
Ment−NorA
Ment−NorB
Ment−Vivi
NorA−NorB
NorA−Vivi
NorB−Vivi
Fst
0.0
0.2
0.4
0.6
0.8
1.0
0.0e+00
5.0e+06
1.0e+07
1.5e+07
2.0e+07
2.5e+07
position (bp)

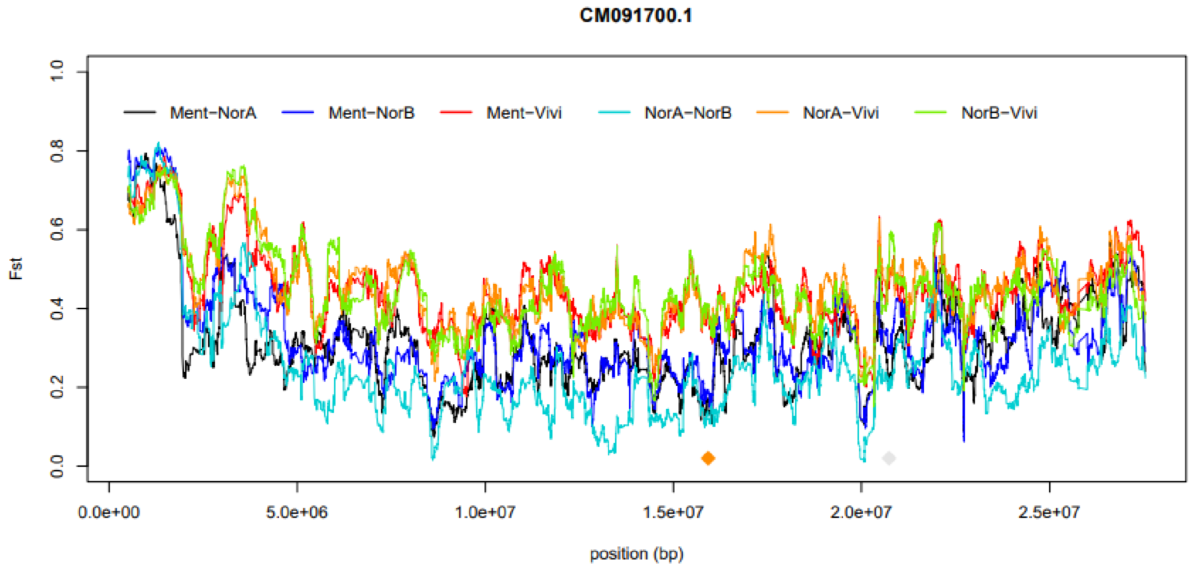
CM091700.1
Ment−NorA
Ment−NorB
Ment−Vivi
NorA−NorB
NorA−Vivi
NorB−Vivi
Fst
0.0
0.2
0.4
0.6
0.8
1.0
0.0e+00
5.0e+06
1.0e+07
1.5e+07
2.0e+07
2.5e+07
position (bp)

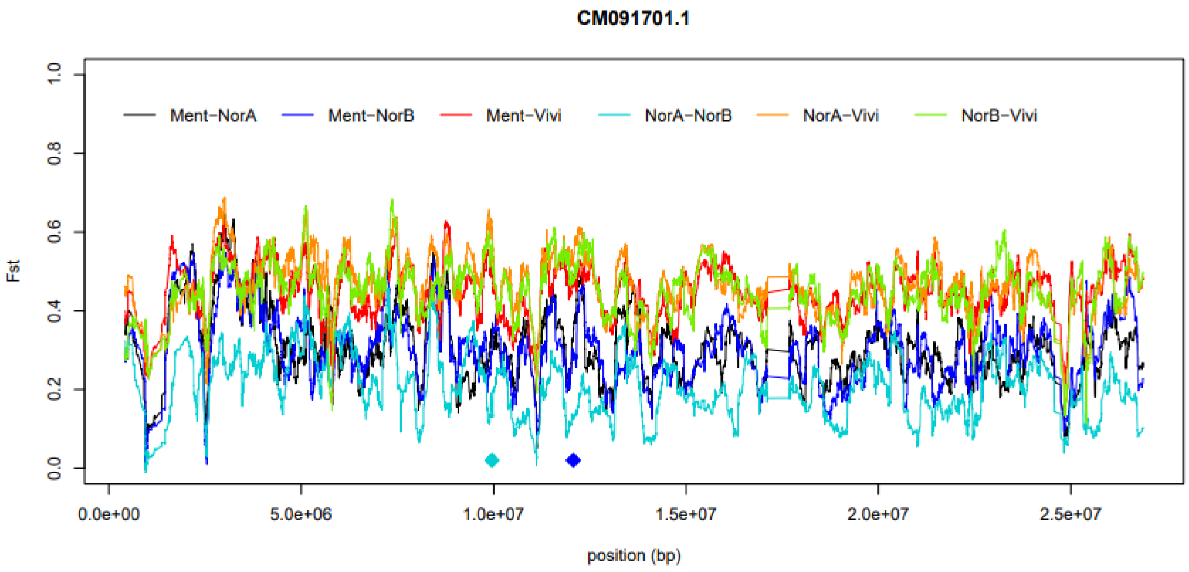
CM091701.1
Ment−NorA
Ment−NorB
Ment−Vivi
NorA−NorB
NorA−Vivi
NorB−Vivi
Fst
0.0
0.2
0.4
0.6
0.8
1.0
0.0e+00
5.0e+06
1.0e+07
1.5e+07
2.0e+07
2.5e+07
position (bp)

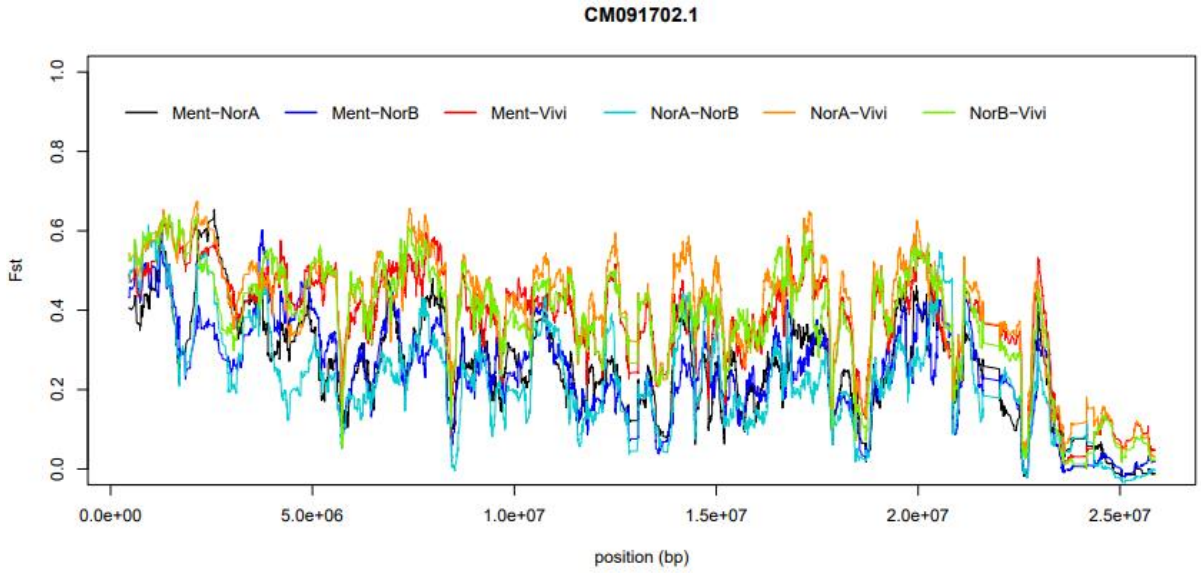
CM091702.1
Ment−NorA
Ment−NorB
Ment−Vivi
NorA−NorB
NorA−Vivi
NorB−Vivi
Fst
0.0
0.2
0.4
0.6
0.8
1.0
0.0e+00
5.0e+06
1.0e+07
1.5e+07
2.0e+07
2.5e+07
position (bp)

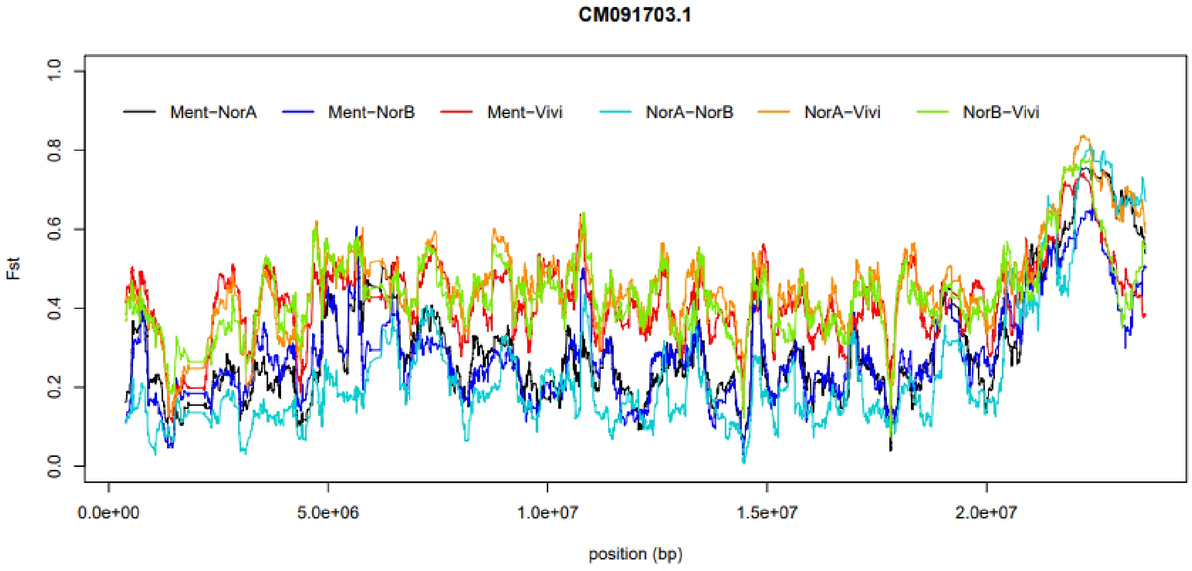
CM091703.1
Ment−NorA
Ment−NorB
Ment−Vivi
NorA−NorB
NorA−Vivi
NorB−Vivi
Fst
0.0
0.2
0.4
0.6
0.8
1.0
0.0e+00
5.0e+06
1.0e+07
1.5e+07
2.0e+07
position (bp)

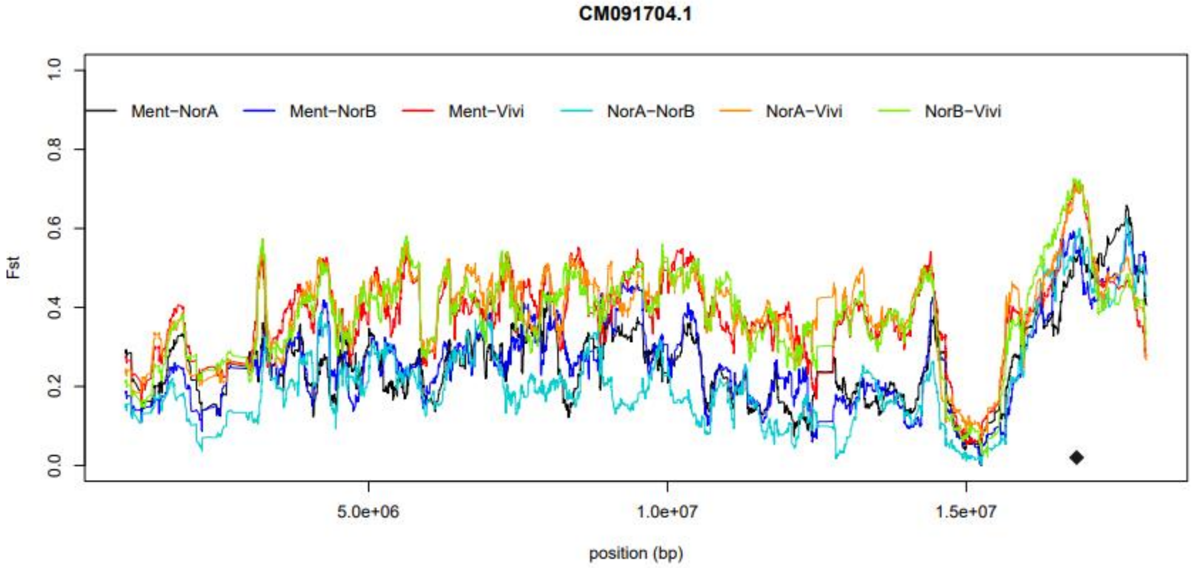

CM091704.1
Ment−NorA
Ment−NorB
Ment−Vivi
NorA−NorB
NorA−Vivi
NorB−Vivi
Fst
0.0
0.2
0.4
0.6
0.8
1.0
5.0e+06
1.0e+07
1.5e+07
position (bp)

**Supplementary Figure 2.** PCA based on 41 SNPs and 2776 fish in *Sebastes* family. Here, reference groups were set as supported by clustering analysis with 56 SNPs (ment = *S. mentella*, viv = *S. viviparus,* norA = *S. norvegicus* A, norB_1/ norB_2 = *S. norvegicus* B subgroups as supported by STRUCTURE analysis). 857 fish analysed only with 41 SNPs are indicated here in yellow ('*unknown*') and were assigned into corresponding clusters using both clustering approaches. Fish of possible hybrid background as indicated by 56 SNPs were not included here.

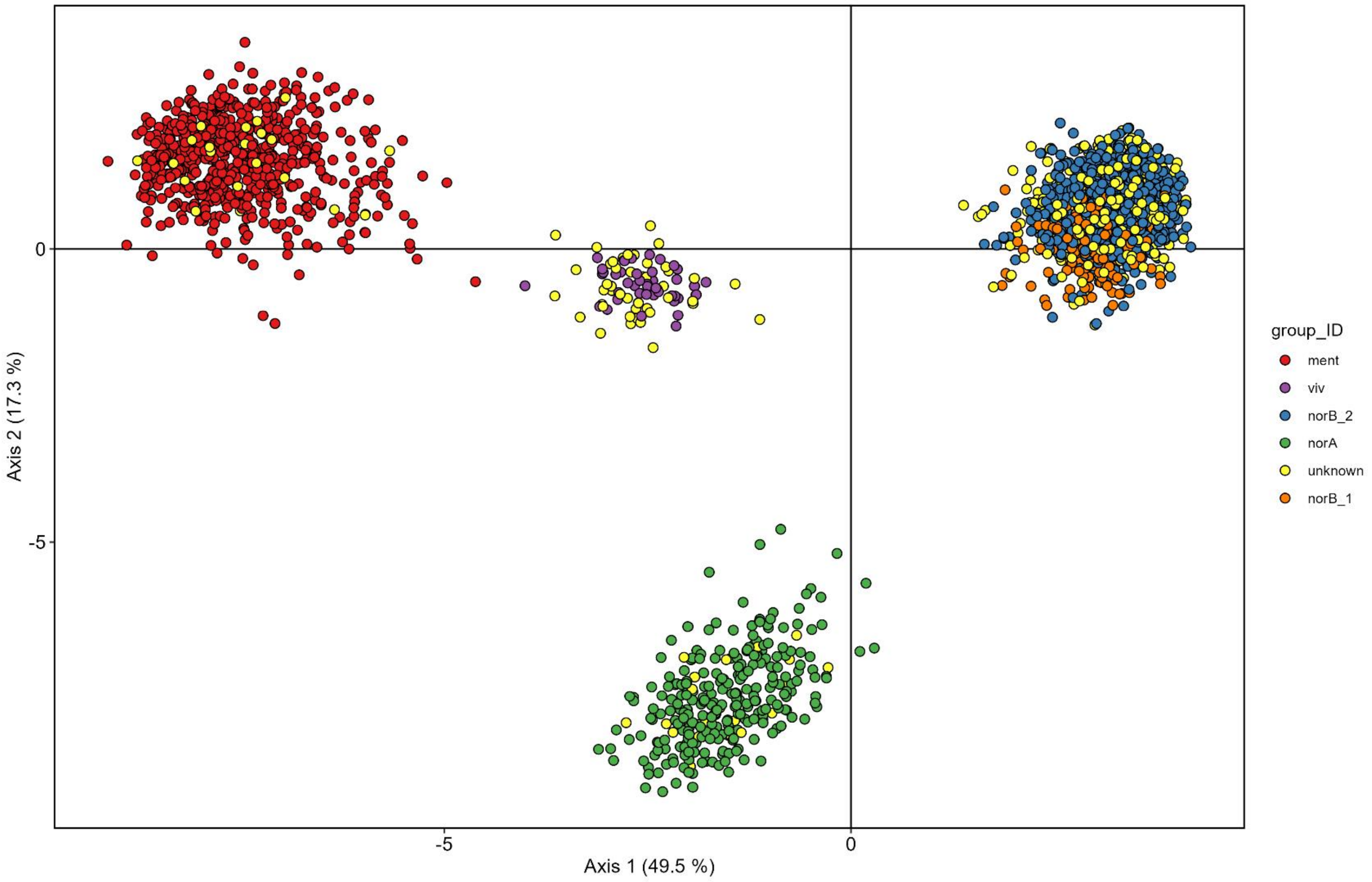

**Supplementary Figure 3.** Partitioning of genetic variation into *K* clusters 2 to 10 as revealed by STRUCTURE analysis. Here, 56 selected SNPs and 1964 fish of *Sebastes* family caught in North-Eastern Atlantic in 2016-2023, were analyzed. Species order follows that shown in Fig.2.

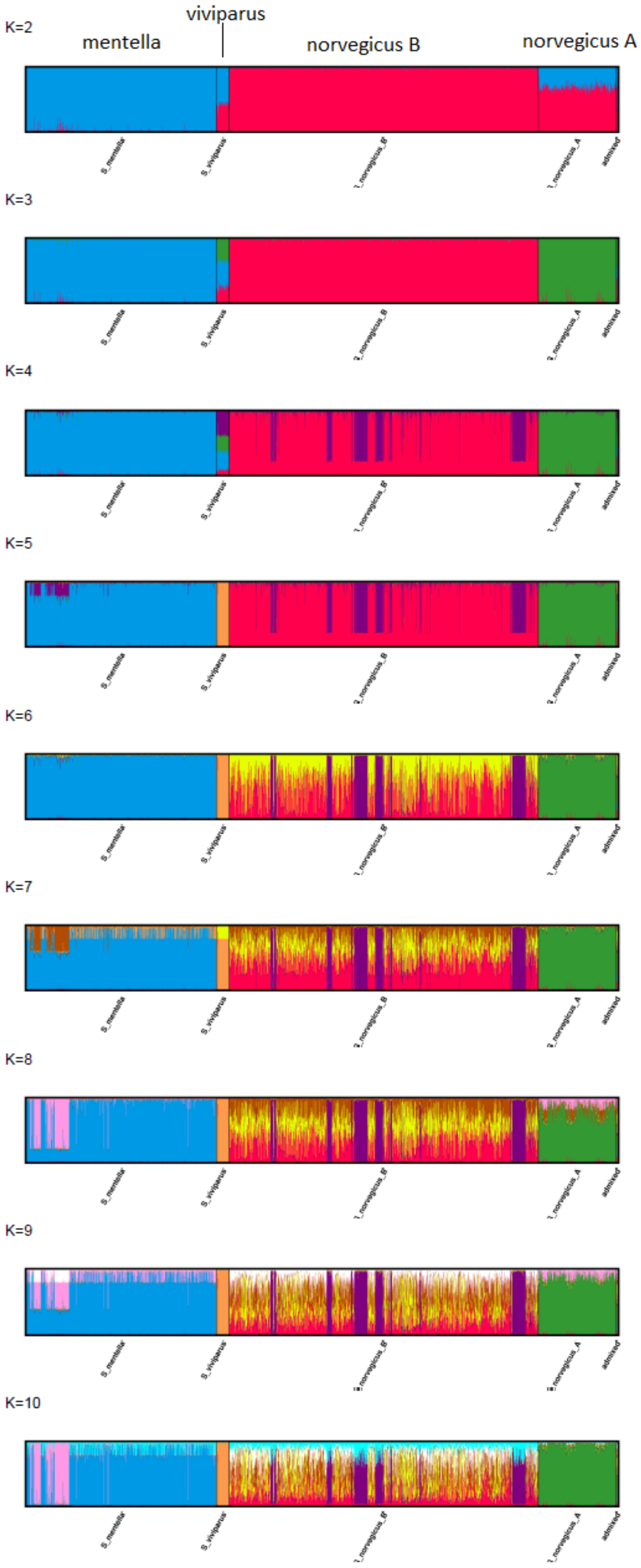

**Supplementary Figure 4.** Probability estimates for different *K* values using different approaches. These results are for the 56 SNPs and 1964 fish. Figure above shows mean Ln Pr(X|K) for different values of *K* that do not show clear peak probability value but plateaus after *K*=5. Second figure from above shows the ad hoc summary statistic for ΔK with a peak value of *K* =2. Four figures below it show summary statistics by Puechmaille.

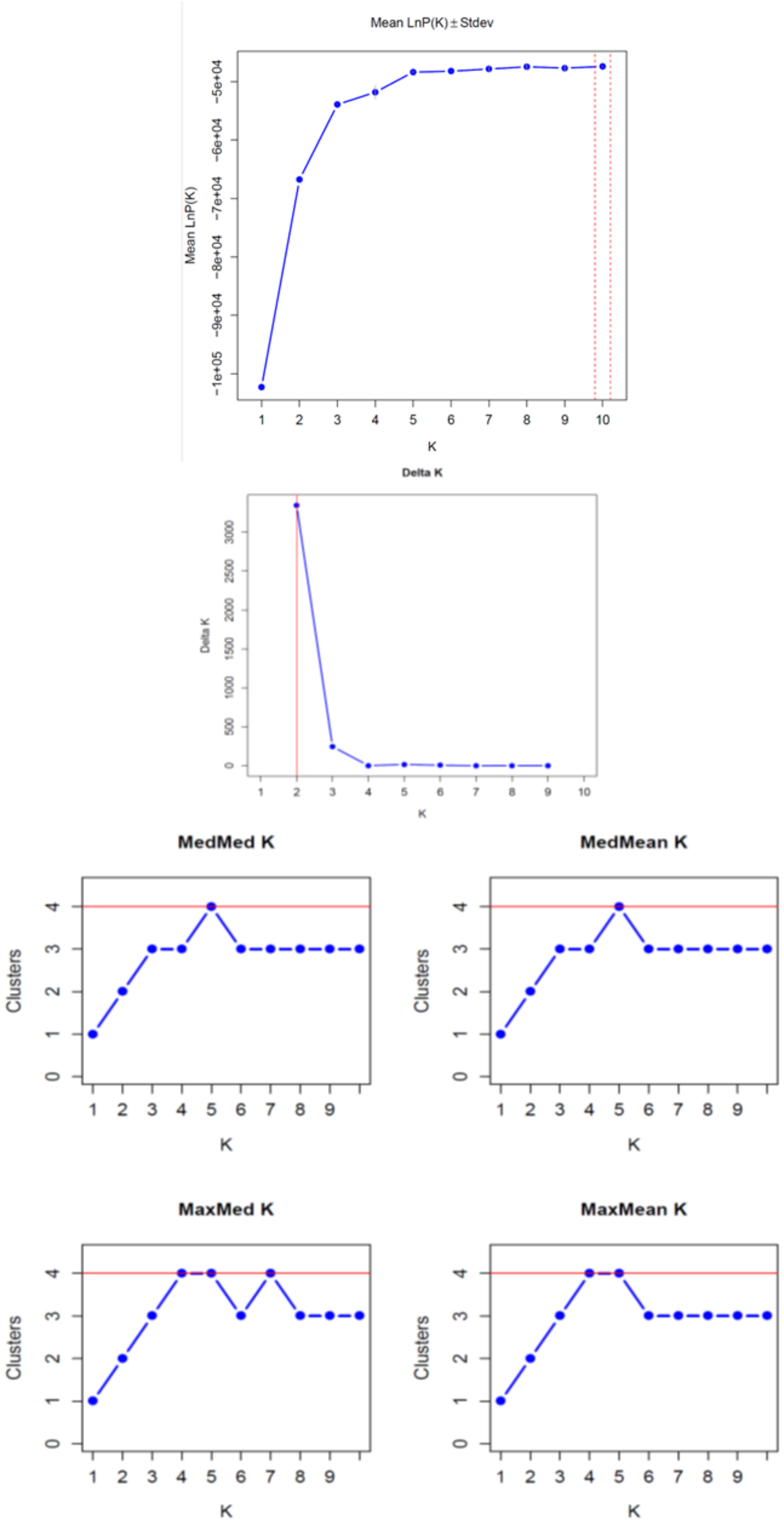

**Supplementary Figure 5**. Partitioning of genetic variation into *K* clusters 2 to 10 as revealed by STRUCTURE analysis. Here, 41 SNPs and 2776 fish were used. Species order follows that shown in Fig.2.

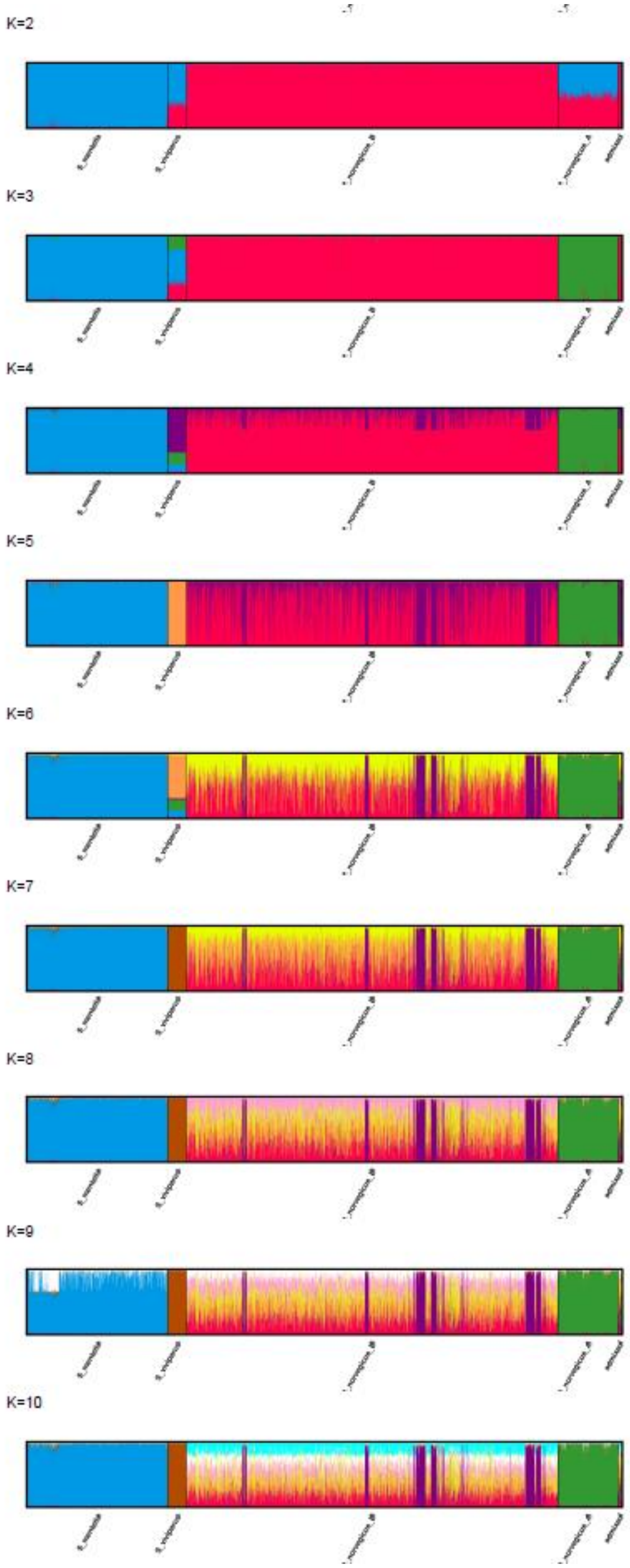

**Supplementary Figure 6.** Probability estimates for different *K* values using 41 SNPs and 2776 fish. Order of figures is the same as in S. Fig 3.

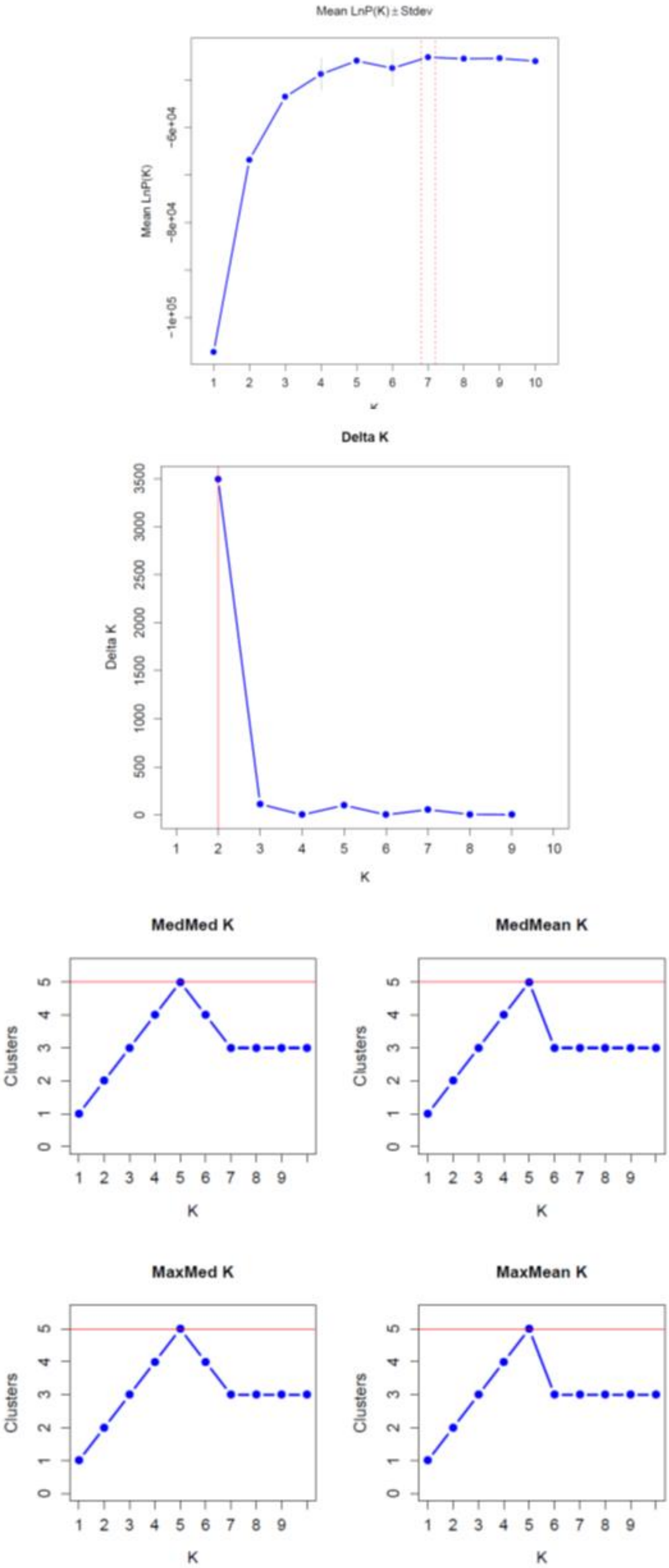

**Supplementary Figure 7. Geographic distribution of species among the collected *Sebastes* samples in this study.**

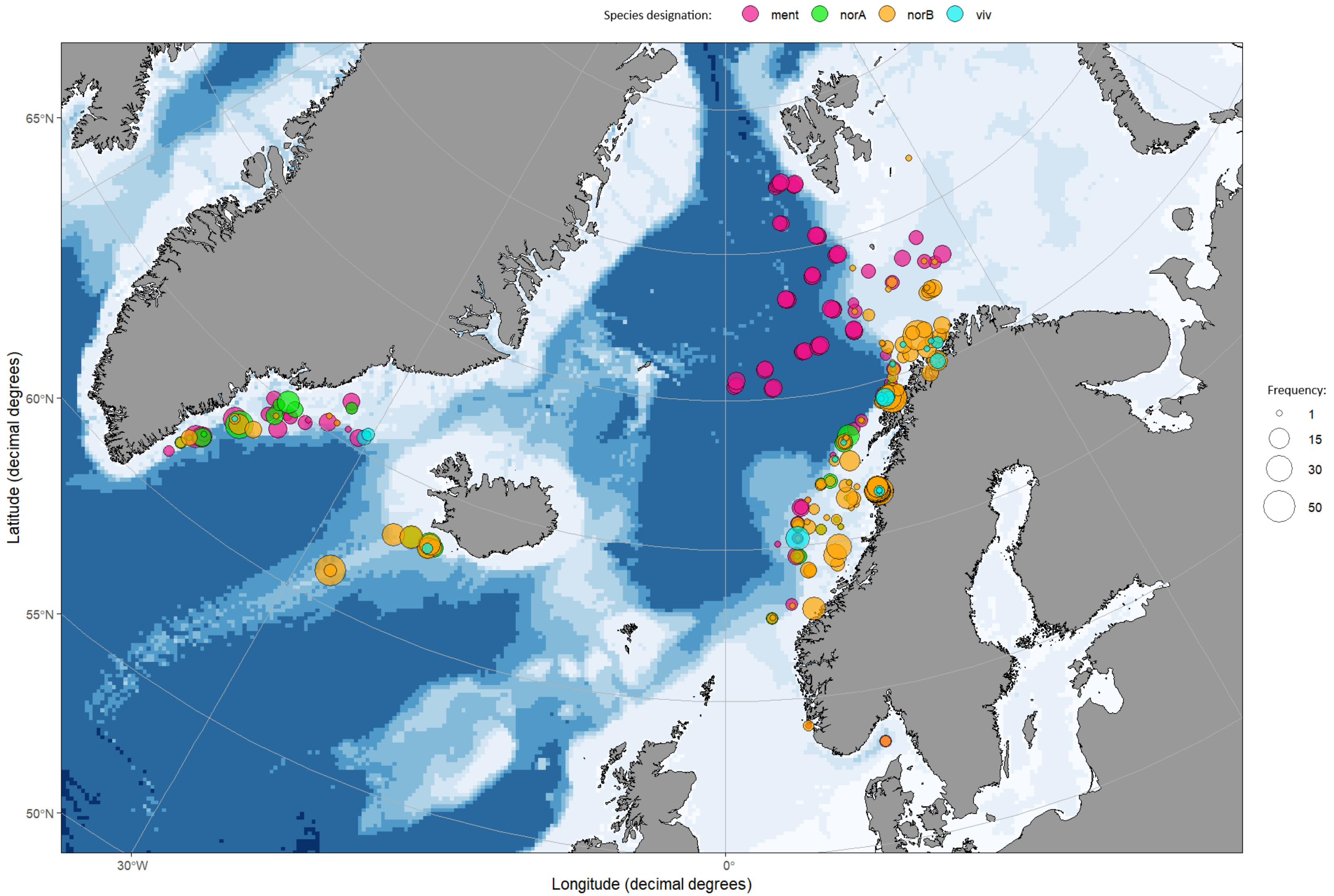

**Supplementary Figure 8. Genetic structure within *S. norvegicus* B as revealed by STRUCTURE clustering analysis with 41 SNPs.** Most supported solutions were $K = 2$-3 as seen from Puechmaille method results below. 177 fish were assigned to giant cluster (shown in green with $K = 3$-5), 742 to the other cluster, and 62 were admixed ($q$ <0.7 to both).

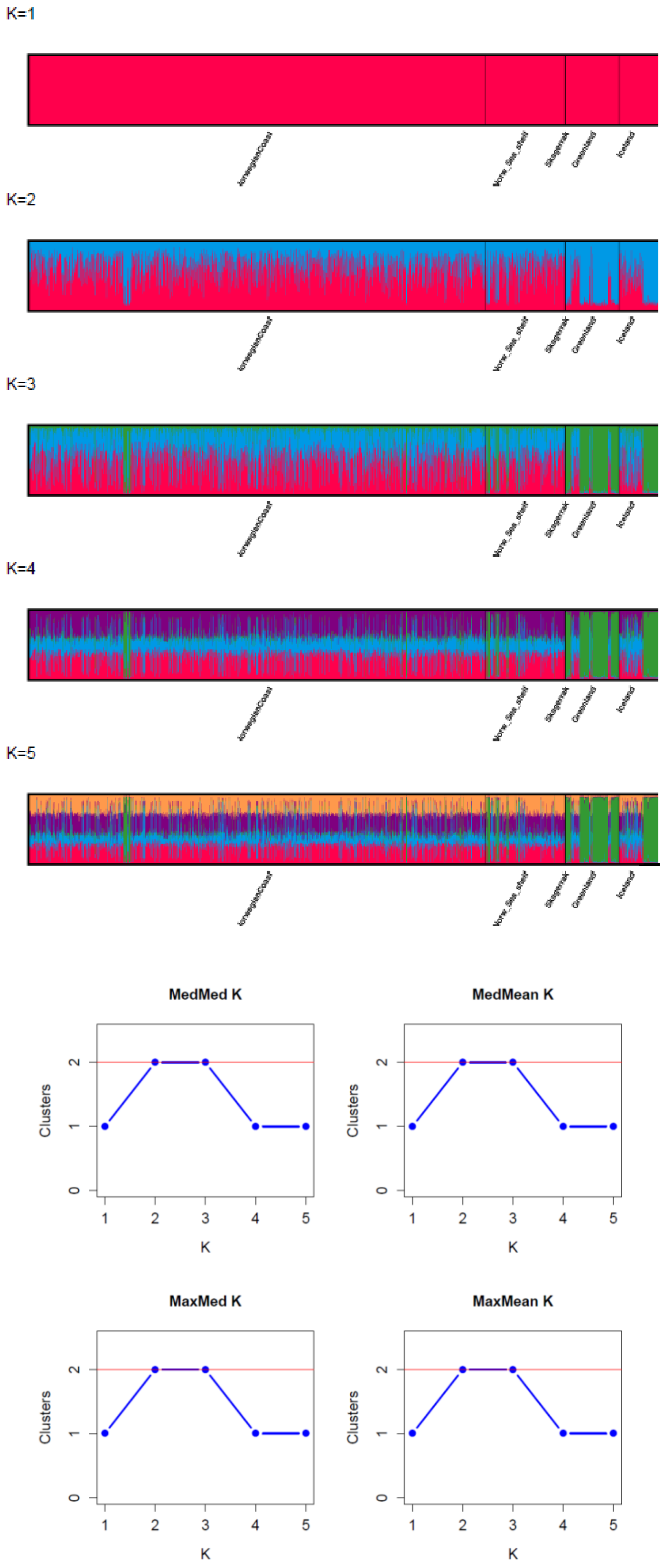

**Supplementary Figure 9. Geographic distribution of *S. norvegicus* B subgroups.** Please note that several samples can be collected from a single position on map. This applies especially to fish assigned into "giant" type in Greenland.

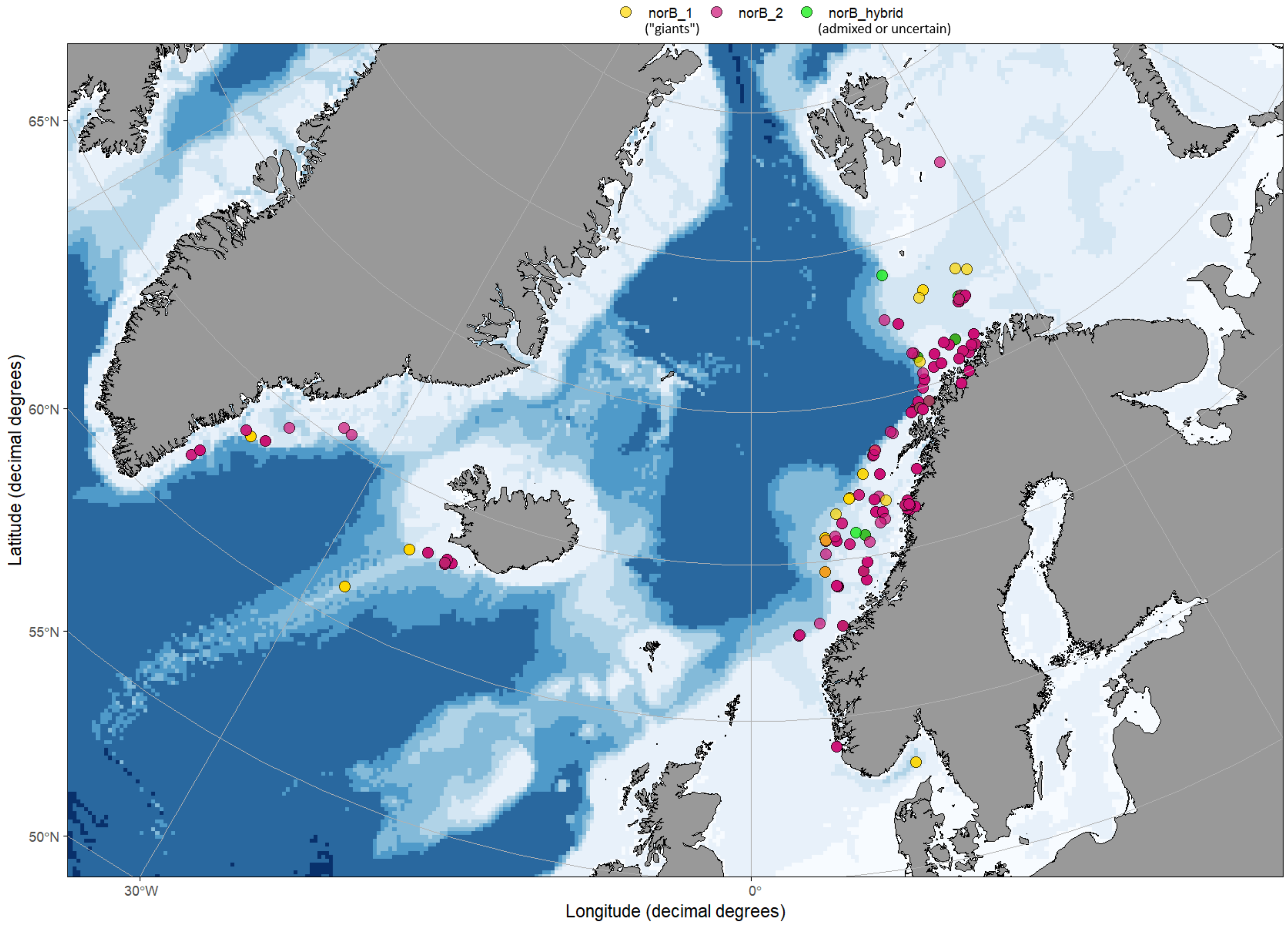

**Supplementary Figure 10.** UPGMA tree of Nei's genetic distances based on 1000 bootstraps. Here, the supported genetic groups within *S. norvegicus* B and *S. mentella* are also shown. Three likely hybrids were removed together with 12 fish of *norvegicus* B and 18 fish in *mentella* subgroups that could not be assigned with high certainty.

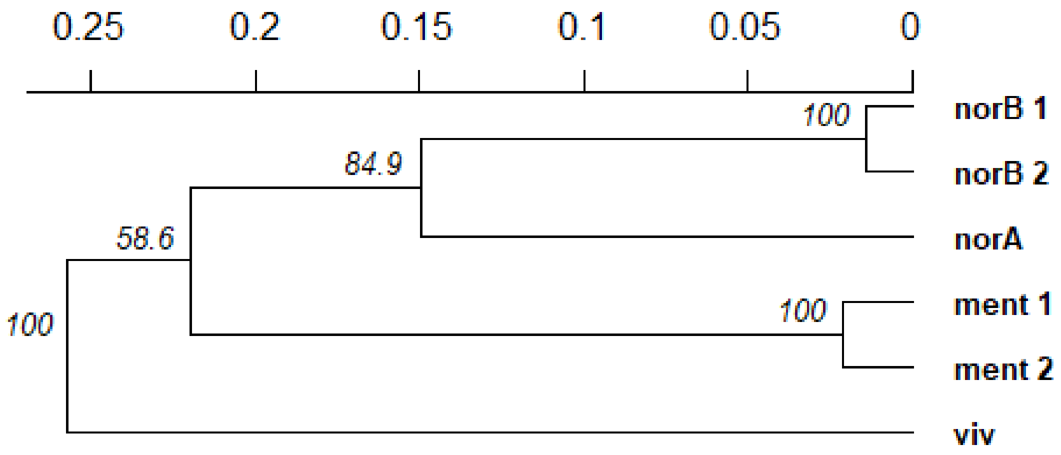

**Supplementary Figure 11a-b. Figure a) shows measured linkage between the used 56 SNP loci in the *S. norvegicus* B dataset (*N* = 1033) as a pairwise matrix.** Linkage was calculated as index of association ($I_A$) with 999 permutation using the *R* package *poppr* (Kamvar, Tabima & Grünwald, 2014). No significant associations were found. Figure b) represents **distribution for the same value in *S. mentella* 56 SNPs dataset (*N* =631**) that didn't show any significant associations either.

**a)**

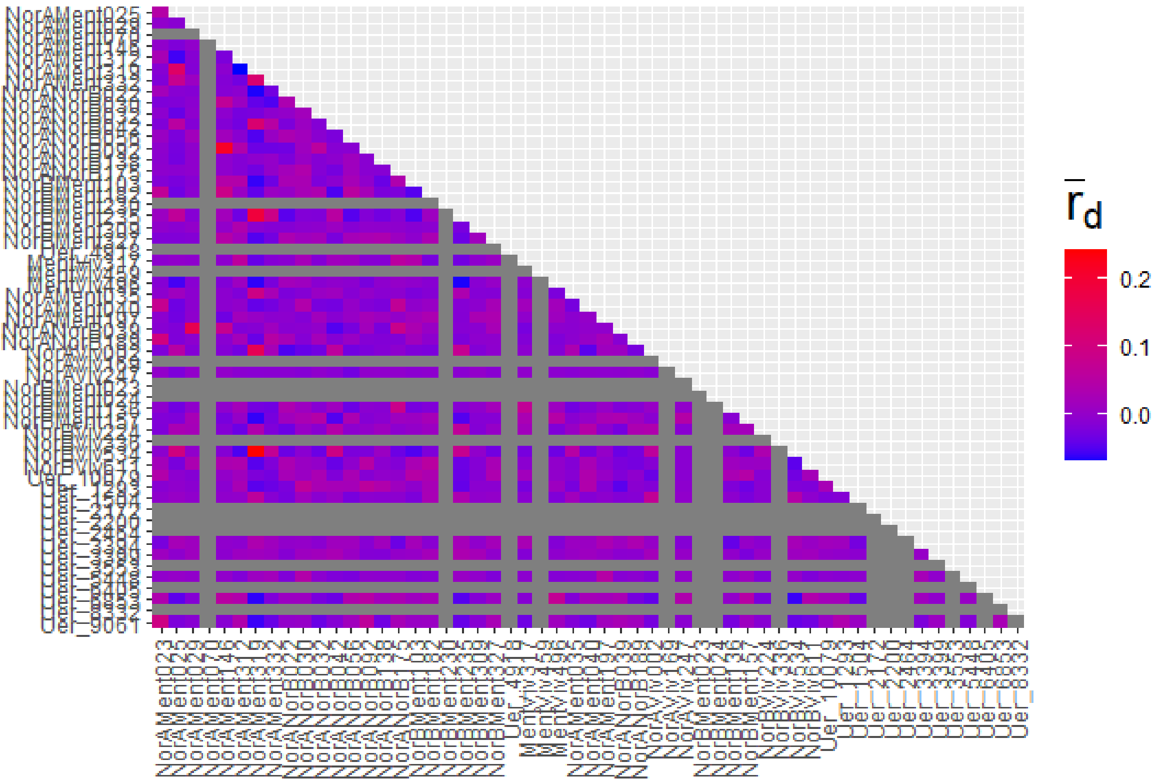


**b)**

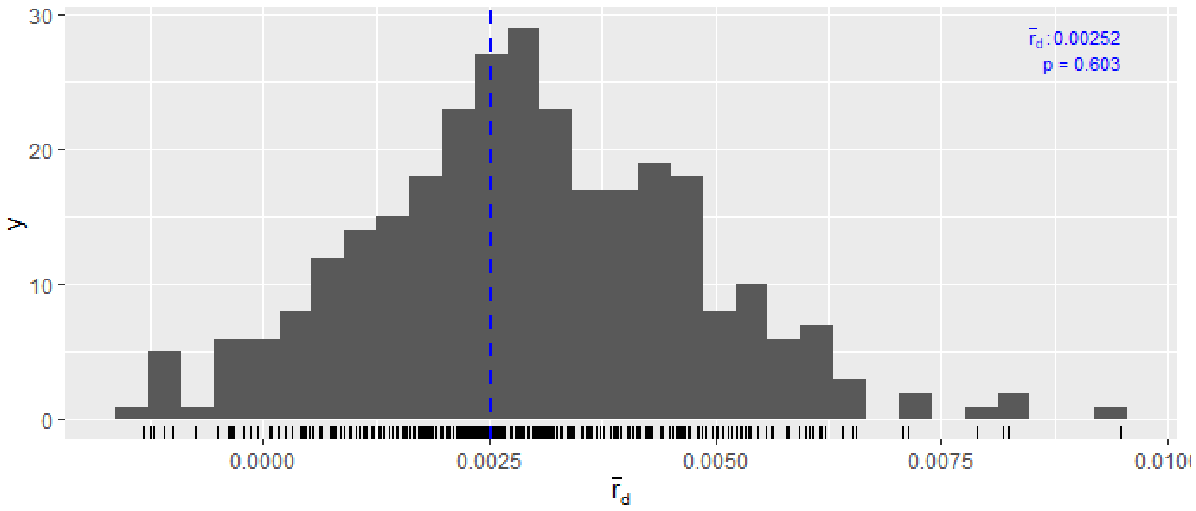

**Supplementary Figure 12. Genetic structure within *S. mentella* as revealed by STRUCTURE clustering analysis with 41 SNPs.** Most supported solutions were $K$ = 2-3 as seen from Puechmaille method results below. 114 fish were assigned to ”Greenlandic” cluster (shown in blue), 466 to the other one, and 49 were admixed ($q$ <0.7 to both).

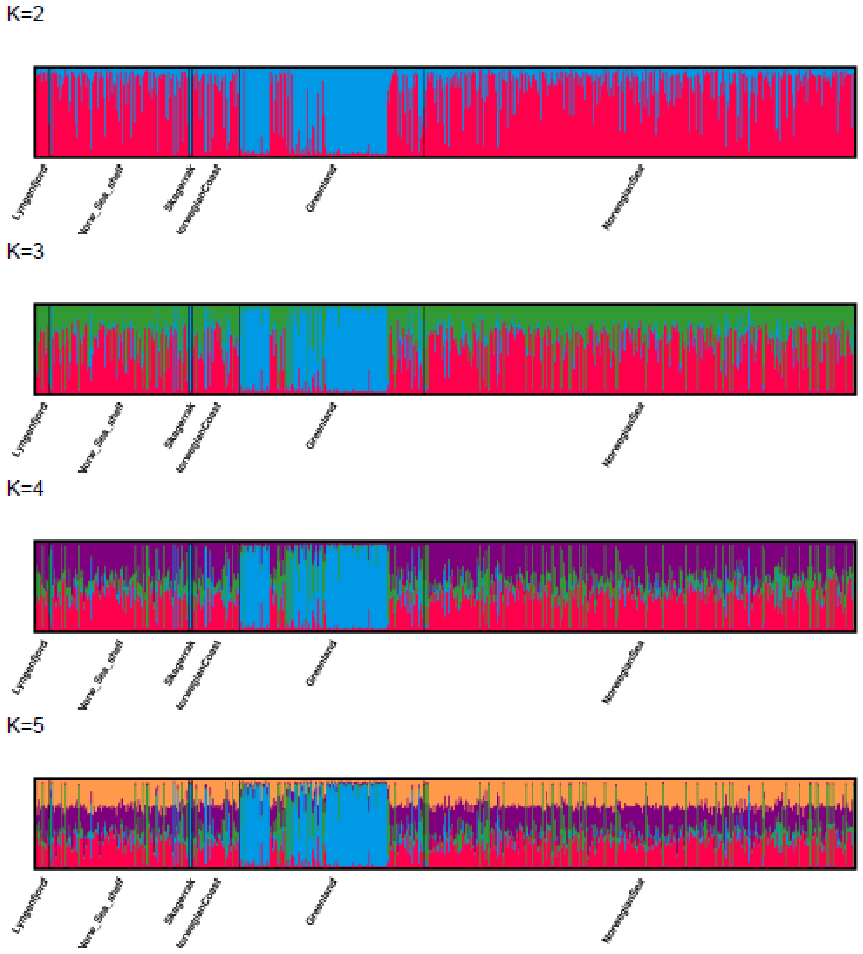


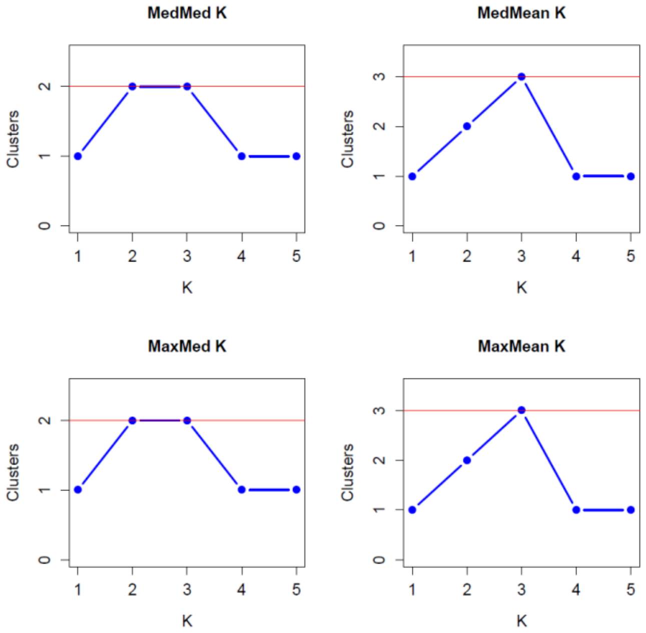

**Supplementary Figure 13a-b). Assignment accuracy**

**A) Assignment accuracy divided on species-level** with different proportions of the 56 SNP loci based on Monte Carlo resampling procedure. Dots indicate outlier observations, whereas mean values are shown with boxplots.

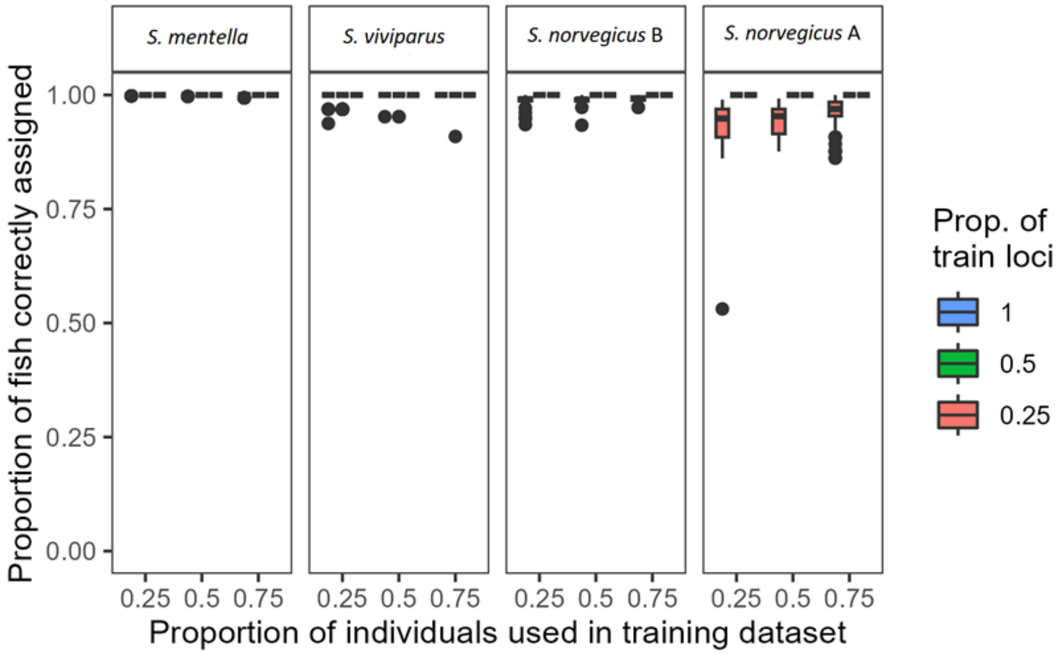


**B) Assignment accuracy with five (sub)species.** Here, *Norvegicus* B was divided into two groups when giants/Greenlandic cluster (number 5 in the figure) was separated from the rest (3). Rest of species are 1 = *S. mentella,* 2 = *S. viviparus*, and 3 = *S. norvegicus* A.

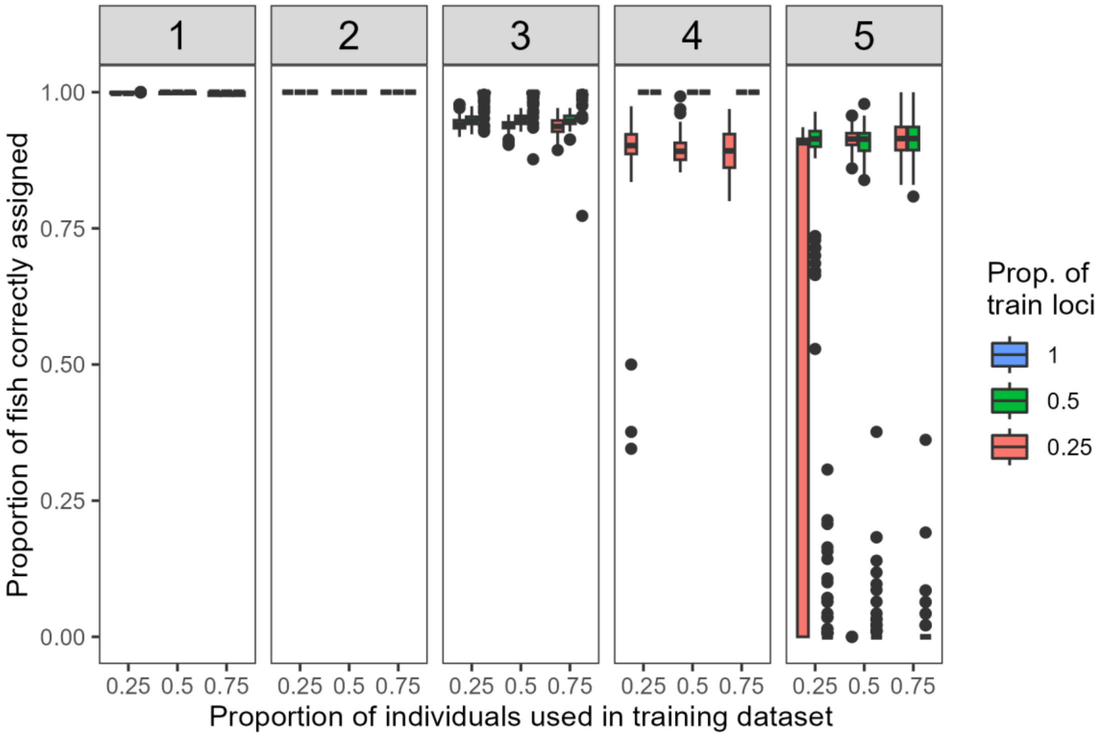

**Supplementary Table 1. Information of the samples included in pool-seq analysis**

| **Species** | **Location** | ***N*** | **year** | **month** | **Lat** | **Lon** | **depth** |
|---|---|---|---|---|---|---|---|
| *S. mentella* | Norway | 15 | 2011 | November | 69.38 | 15.14 | 575 |
| | Norway | 5 | 2016 | NA | 68.16 | -3.94 | 355-615 |
| *S. norvegicus* B | Norway | 4 | 2001 | October | 69.33 | 15.13 | 324 |
| | Norway | 20 | 2002 | October | 68.20 | 11.15 | 195 |
| | Greenland | 20 | 2011 | August | 62.20 | 40.65 | 430 |
| *S. norvegicus* A | Greenland | 9 | 2011 | NA | 64.26 | 25.15 | 370 |
| | Norway | 5 | 2014 | NA | 65.66 | 5.86 | 354-372 |
| | Norway | 2 | 2001 | November | 69.33 | 15.13 | 324 |
| *S. viviparus* | Faroe Island | 19 | 2002 | September | 60.91 | -6.07 | 375-426 |

**Supplementary Table 2. Information of the samples included in SNP analysis**

| Larger area | Year | Population ID | Number of fish collected |
|---|---|---|---|
| Greenland | 2012 | Gre_12 | 53 |
| | 2019 | Gre_19 | 66 |
| | 2020 | Gre_20 | 76 |
| | 2022 | Gre_22 | 246 |
| Iceland | 2017 | Ice_17 | 143 |
| Norwegian Sea | 2019 | Nor_sea_19 | 344 |
| Norwegian shelf | 2016 | ref_shelf_16 | 186 |
| | 2018 | ref_shelf_18 | 54 |
| | 2019 | ref_shelf_19 | 105 |
| | 2020 | ref_shelf_20 | 51 |
| | 2021/2022 | ref_shelf_21_22 | 15 |
| Norwegian coast | 2017 | ref_coast_17 | 100 |
| | 2019 | ref_coast_19 | 120 |
| | 2020 | ref_coast_20 | 346 |
| | 2021 | ref_coast_21 | 326 |
| | 2022 | ref_coast_22 | 569 |
| | 2023 | ref_coast_23 | 97 |
| Lyngenfjord | 2020 | Lyn_coast_20 | 12 |
| Skagerrak | 2023 | Ska_23 | 5 |
| total | | | 2914 |

**Supplementary Table 3. Geographic distribution of *Sebastes* species based on *dapc* clustering analysis with the selected 56 SNP and highest species-level clustering (*K*=4).** Numbers with dark grey background indicate most common species in that specific sample whereas light grey background indicates less common species in that sample, and white background that the species was not observed in this sample.

| **Area** | site_year | *norvegicus A* | *norvegicus B* | *mentella* | *viviparus* |
|---|---|---|---|---|---|
| **Greenland** | Gre_19 | 36 | 29 | 0 | 0 |
| | Gre_20 | 0 | 38 | 1 | 0 |
| | Gre_22 | 63 | 25 | 147 | 11 |
| | Gre_giant_12 | 0 | 24 | 0 | 0 |
| **Iceland** | Ice_17 | 69 | 70 | 0 | 2 |
| **Norwegian shelf** | ref_shelf_16 | 29 | 45 | 104 | 1 |
| | ref_shelf_18 | 5 | 45 | 3 | 0 |
| | ref_shelf_19 | 0 | 91 | 0 | 14 |
| | ref_shelf_20 | 7 | 41 | 2 | 0 |
| | ref_shelf_21_22 | 0 | 10 | 3 | 2 |
| **Norwegian Sea** | Norw_sea_19 | 0 | 0 | 344 | 0 |
| **Lyngenfjord** | Lyn_coast_20 | 0 | 0 | 11 | 1 |
| **Norwegian coast** | ref_coast_17 | 0 | 98 | 0 | 0 |
| | ref_coast_19 | 19 | 38 | 6 | 10 |
| | ref_coast_20 | 2 | 84 | 3 | 0 |
| | ref_coast_21 | 15 | 173 | 0 | 1 |
| | ref_coast_22 | 14 | 122 | 4 | 1 |
| | ref_coast_23 | 0 | 96 | 0 | 0 |
| **Skagerrak** | Ska_23 | 0 | 2 | 3 | 0 |
| | | 259 | 1031 | 631 | 43 |

**Supplementary Table 4. Species-level diversity indices measured with 56 SNPs (above) and 41 SNPs (below).**

Species membership was defined based on two clustering approaches. Likely hybrids were removed.

Explanation for each variable is given below

***56 SNPs dataset***

| Species | *N* | *Hexp* | *Hobs* | *Fis* | *Fis_95%_ll* | *Fis_95%_ul* | *non-HWE prop* |
|---|---|---|---|---|---|---|---|
| *S. mentella* | 634 | 0.202 | 0.183 | 0.082 | 0.047 | 0.166 | 0.196 |
| *S. viviparus* | 47 | 0.073 | 0.069 | 0.298 | -0.033 | 0.248 | 0 |
| *S. norvegicus_A* | 260 | 0.124 | 0.122 | 0.028 | -0.009 | 0.056 | 0 |
| *S. norvegicus_B* | 1033 | 0.109 | 0.099 | 0.098 | 0.038 | 0.173 | 0.107 |

***41 SNPs dataset***

| Species | *N* | *Hexp* | *Hobs* | *Fis* | *Fis_95%_ll* | *Fis_95%_ul* | *non-HWE prop* |
|---|---|---|---|---|---|---|---|
| *S. mentella* | 655 | 0.171 | 0.157 | 0.072 | 0.032 | 0.157 | 0.195 |
| *S. viviparus* | 88 | 0.082 | 0.085 | -0.045 | -0.151 | 0.052 | 0.024 |
| *S. norvegicus_A* | 282 | 0.137 | 0.131 | 0.046 | 0.013 | 0.078 | 0.024 |
| *S. norvegicus_B* | 1742 | 0.107 | 0.097 | 0.098 | 0.033 | 0.168 | 0.171 |

N number of fish, Hexp expected heterozygosity, Hobs observed heterozygosity, Fis inbreeding coefficient, Fis_95%_ll/hl lower and upper 95% confidence interval based on 1999 bootstraps, non-HWE prop. Proportion of loci not in HWE after FDR correction

**Supplementary Table 4. Pairwise genetic divergence measured as pairwise *F*st.**
Upper left corner show results for 41 SNPs and lower right corner for 56 SNPs.
Statistical significance was tested with 1999 bootstraps to obtain corresponding p-values.
All p-values were zero.

| | *S. mentella* | *S. viviparus* | *S. norvegicus* B | *S. norvegicus* A |
|---|---|---|---|---|
| *S. mentella* | | 0.726 | 0.749 | 0.714 |
| *S. viviparus* | 0.641 | | 0.789 | 0.789 |
| *S. norvegicus* B | 0.677 | 0.757 | | 0.719 |
| *S. norvegicus* A | 0.633 | 0.772 | 0.669 | |
